# Spatial resolution of an integrated C_4_+CAM photosynthetic metabolism

**DOI:** 10.1101/2021.11.25.470062

**Authors:** Jose J Moreno-Villena, Haoran Zhou, Ian S Gilman, S. Lori Tausta, C. Y. Maurice Cheung, Erika J Edwards

## Abstract

C_4_ and CAM photosynthesis have repeatedly evolved in plants over the past 30 million years. Because both repurpose the same set of enzymes but differ in their spatial and temporal deployment, they have long been considered as distinct and incompatible adaptations. Remarkably, *Portulaca* contains multiple C_4_ species that perform CAM when droughted. Spatially explicit analyses of gene expression reveal that C_4_ and CAM systems are completely integrated in *P. oleracea*, with CAM and C_4_ carbon fixation occurring in the same cells and CAM-generated metabolites likely incorporated directly into the C_4_ cycle. Flux balance analysis corroborates the gene expression and predicts an integrated C_4_+CAM system under drought. This first spatially explicit description of a C_4_+CAM photosynthetic metabolism presents a new blueprint for crop improvement.

## Introduction

C_4_ and CAM photosynthesis are two important adaptations that have evolved multiple times in terrestrial plants (Edwards, 2019). These adaptations allow plants to maintain or increase their autotrophic core carbon metabolism in adverse conditions. Both act as carbon concentration mechanisms (CCMs) that alleviate energy losses caused by photorespiration, which can occur when atmospheric CO_2_ levels are low, internal leaf temperatures are high, or plant stomatal conductance is reduced, for instance, due to water stress (Sage, *et al*., 2012).

Both CCMs have repurposed a shared set of core metabolic enzymes that are present in all plants but differ in how they isolate and create a CO2-enriched environment around Rubisco, the enzyme that fixes atmospheric CO_2_ into sugars via the Calvin cycle. In C_4_ metabolism, PEP carboxylase (PEPC) first interacts with dissolved CO_2_ in mesophyll cells to form a temporary 4-carbon molecule, typically malate or aspartate. These molecules are then transported to inner bundle sheath cells, where Rubisco is confined, to be decarboxylated and release CO_2_ at high rates that saturate Rubisco. There, CO_2_ enters the Calvin cycle and is assimilated into carbohydrates (Hatch and Slack, 1966). Thus, C_4_ is essentially a temporally synchronous two-cell photosynthetic system, with separate compartments for PEPC and Rubisco that results in C_4_ plants achieving the highest rates of photosynthesis (Black, Chen and Brown, 1969). In CAM plants, stomata open and CO_2_ fixation by PEPC occurs during the night. The 4-carbon malate is stored overnight in the form of malic acid in mesophyll cell vacuoles. During the day stomata close, malate exits the vacuole for decarboxylation, and CO_2_ is released and assimilated by the Calvin cycle in the same cell (Osmond, 1978) .Thus CAM is a temporally asynchronous, single cell photosynthetic system, with initial carbon capture by PEPC occurring at night, but eventual Rubisco assimilation and carbohydrate production occurring during the day in the same cell. The CAM inverted stomatal behaviour provides increased water use efficiency by avoiding water loss through stomata during the hottest daytime hours (Winter, Aranda and Holtum, 2005). There is also significant variability in the degree to which CAM is expressed: many species use CAM as their primary metabolism, but possibly more common is a facultative CAM system, where plants operate a C3 metabolism but exhibit facultative CAM as a stress response (“C3+CAM” sensu Edwards 2019). Both CCMs exhibit remarkable evolutionary convergence. C_4_ has evolved at least 60 times, and C_4_ species include some of the most productive plants on Earth, comprising upwards of 30% of gross terrestrial primary productivity and including several essential crops such as maize and sorghum (Sage, Christin and Edwards, 2011). While the number of CAM origins is less well known, it is speculated to be higher than C_4_, and some form of CAM metabolism is dominant in a variety of ecosystems (Silvera *et al*., 2009).

Despite the large number of independent origins of both C_4_ and CAM, for the most part plant lineages tend to evolve one CCM or the other. Over 40 years ago, *Portulaca oleracea* was identified as the first known C_4_ plant that also operates a facultative CAM cycle (C_4_+CAM) in response to drought or photoperiod (Koch and Kennedy, 1980). Full integration of C_4_ and CAM cycles, whereby both operate in the same population of photosynthetic mesophyll cells, seemed implausible on multiple fronts: 1) the shared set of enzymes would need to be expressed at different times of day and in different cells, creating significant regulatory conflict; 2) the activity of Rubisco (for CAM) and PEPC (for C_4_) in the same mesophyll cells during the day would result in futile competition for CO_2_ and metabolic cycling; 3) the anatomical requirements for optimizing each CCM are distinct and potentially antagonistic, in that CAM requires large mesophyll cell volume yet C_4_ requires a high bundle sheath to mesophyll volumetric ratio; 4) the adaptations themselves are potentially ecologically antagonistic, as the benefits to CAM are primarily to increase water use efficiency, whereas C_4_ plants take advantage of high light and seasonally available water to achieve fast growth and high productivity (Sage, 2002). Only a handful of immunolocalization studies have pursued the spatial resolution of C_4_+CAM in *Portulaca* (Guralnick *et al*., 2002; Lara, Drincovich and Andreo, 2004) and results were equivocal due to lack of C4-specific vs CAM-specific molecular markers. Regardless, the most accepted hypothesis has been that CAM and C_4_ cycles operate independently in *Portulaca*, with C_4_ in bundle sheath and mesophyll cells and CAM in specialized water storage cells. Only (Lara, Drincovich and Andreo, 2004) suggested a possible integration-that malate generated from CAM could theoretically be processed in the C_4_ bundle sheath.

Recent genomic information for *Portulaca* has provided an opportunity to resolve the spatial configuration of C_4_ and CAM reactions. Multiple studies have now identified C_4_ and CAM-specific homologs of PEPC and some accompanying CCM-affiliated enzymes (Christin *et al*., 2014; Ferrari *et al*., 2020; Gilman *et al*., 2021), providing key markers to identify the cell populations where CAM and C_4_ CO_2_ fixation occur. The lack of significant upregulation of distinct homologs of CAM decarboxylation genes in multiple *Portulaca* species is notable (Gilman *et al*., 2021), and suggests that some elements of C_4_ and CAM biochemistry may be shared. However, these analyses were performed on whole leaf transcriptomes and lacked the spatial resolution needed to identify where C_4_ and CAM processes were occurring. We performed two different spatially explicit gene expression analyses of *Portulaca oleracea* leaves under well-watered and drought conditions that isolated mesophyll and bundle sheath cell populations. Moreover, we utilized constraint-based modeling to specifically test for the most efficient biochemical model of C_4_ and CAM integration across mesophyll and bundle sheath cells under a variety of scenarios. Together, we present these analyses of *Portulaca* as a first description of a novel plant photosynthetic metabolism.

## Results

### Assessment of CAM induction

Titratable acidity analysis of well-watered *P. oleracea* leaves at 7h and 19h did not result in significant accumulation of acids overnight (two-sample t-test, p = 0.62, t = -0.33, df = 5.95; Fig.1-A, Table S1). This confirmed little or no accumulation of malic acid from CAM activity under well-watered conditions. After seven days of total water withholding, significant accumulation of acids was detected in leaves collected early in the morning compared to those collected at the end of the photoperiod (t-test, p = 0.00032, t = 7.34, df = 5.17; Fig. 1-A, Table S1), confirming CAM induction.

**Fig. 1.**
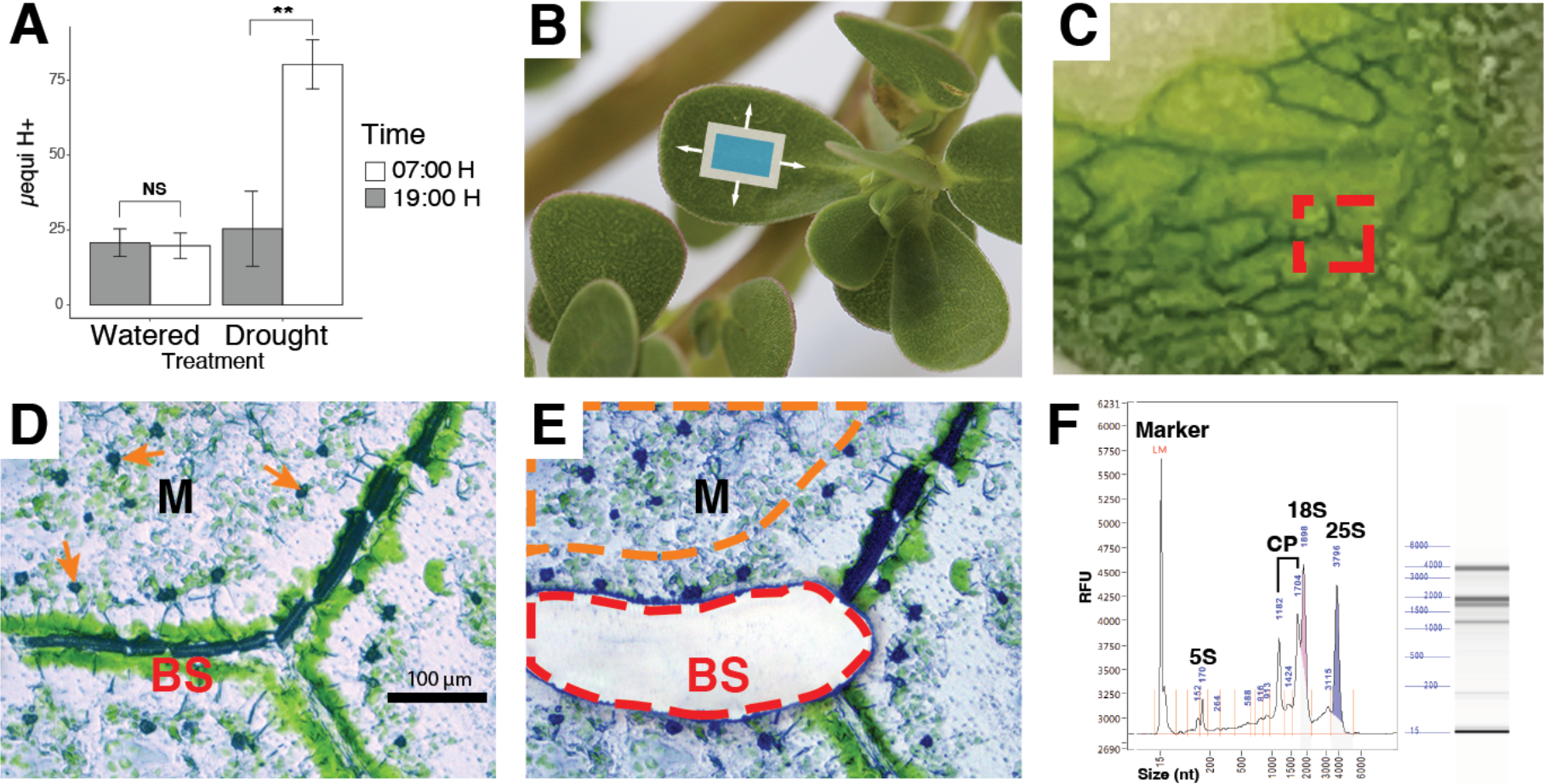
Drought induction of CAM and laser capture microdissection. (A) Diel fluctuation of titratable acidity from whole leaves of well-watered plants and after a 7-day drought treatment. (B) *Portulaca oleracea*, illustrating the orientation of paradermal sections (blue box). (C) Fresh, flash-frozen paradermal leaf section in cryosection block; red square indicates an area used for tissue dissection. (D) Microphotograph of a 12um thick leaf paradermal section indicating bundle sheath (BS) and mesophyll (M) tissues before (D), and after (E), BS cell capture. Orange arrows in (D) indicate calcium oxalate crystals in M cells. The red line in (E) indicates a laser-cut area of BS tissue, and the orange line illustrates an area of M tissue for laser capture. (F) RNA profile by electrophoresis for quality control. CP: Chloroplast RNAs.

### LMD-RNA Sequencing and read alignments

To obtain cell-specific gene expression, we made paradermal sections from well-watered and droughted *Portulaca oleracea* leaves (Fig. 1A-B; Table S1) and captured groups of mesophyll and bundle sheath cells using laser microdissection (LCM) (Fig. 1, C to E). We isolated mRNA from populations of each tissue for short read sequencing. We sequenced 28 libraries that represented 1.05 billion 100 bp paired-end reads with a mean of 37.49 million (SD = 3.85 million) reads per library (Figure 2; Table S2; Methods). After quality filtering and trimming, a mean of 68.97% of the reads were kept. The *P. oleracea* Trinity assembled transcriptome retrieved from Gilman et al. (2021) consisted of 444307 contigs and 230895 Trinity “unigenes”. Transdecoder predicted coding sequences in 152530 of the contigs and CDHIT reduced redundant contigs to a final set of 54241 contigs in 37518 “unigenes” (Online Table S1). The reduced dataset of mRNA contigs represented 83.20 Mbp with a 2101 bp N50. The mean reads mapped to the reduced set of *P. oleracea* mRNA contigs by Kallisto was 75.30% per library (SD = 3.65%), and a mean of 59.96% (SD= 3.48%) of mapped reads had unique alignments (Supplement Table S2). Reads mapping to CCM and starch metabolism related gene families constituted 10.97% (SD = 2.49%) of daytime mesophyll cell libraries and 12.18% (SD = 2.94%) of the daytime bundle sheath cell libraries. On average, 5.20% (SD = 1.83%) of reads from nocturnal libraries mapped to CCM-related genes, while 11.62% (SD=2.70%) of reads mapped to CCM-related genes in daytime libraries.

**Fig. 2.**
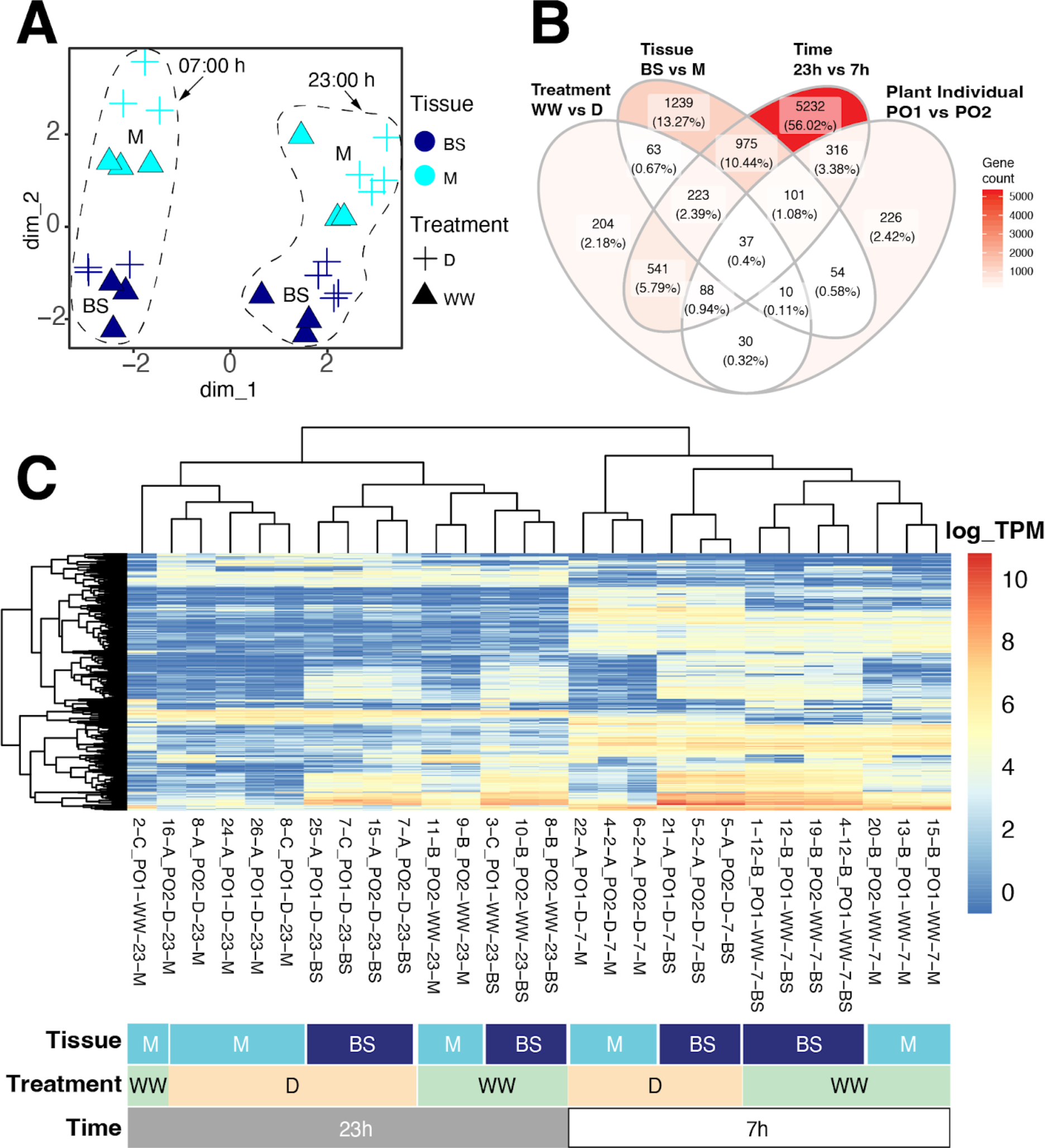
Transcriptome-wide gene expression patterns across mesophyll and bundle sheath samples. (A) All LMD samples projected into the first two dimensions of a transcriptome-wide multidimensional scaling analysis of log-transformed expression. Treatments: D, drought. WW, well-watered. (B) Venn diagram with number of differentially expressed genes across variables. (C) Heatmap with the log of gene expression of the 500 most variable genes across samples. Dendrograms cluster mRNA libraries (on the top) and genes (on the left) based on gene expression similarities.

### Global transcriptional differences across experimental variables

Multidimensional scaling of transcriptome-wide gene expression clustered time points and tissue replicates (n >= 3) along dimensions 1 and 2, respectively (Fig. 2A). MDS further separated samples by water status within each time and tissue cluster. Of the 22509 unigenes that passed the low read mapping filter, 2702 (12.00%) were differentially expressed (DE) between M and BS (padj < 0.05; Fig. 2B; online Table S2). Of those, 1030 (38.12%) were more highly expressed in M and 1672 (61.88%) in BS. 1196 unigenes (5.31%) were significantly DE across watering treatments; of those, 745 (62.29%) were more abundant in well-watered plants and 451 (37.71%) were more abundant during drought. We found 7513 DE unigenes (33.37%) across time points, with 3553 (47.29%) and 3960 (52.71%) more abundant during the day and night, respectively.

Using all samples, gene differential expression analyses revealed that most detected gene expression shifts occurred between day and night, followed by cell type and water status (Fig. 2B). Gene ontology (GO) analyses of differentially expressed genes showed mesophyll enrichment of non-photosynthetic carbon capturing GO terms (carbon utilization GO:0015976) while Calvin cycle (GO:0019253) and photorespiratory terms (GO:0019264,GO:0009853) were enriched in bundle sheath cells (Fig. S1). Genes upregulated under drought were enriched in stress response GO terms as response to salt and abscisic acid (GO:0009651;GO:0009737), while genes more highly expressed when well-watered were enriched in photosynthesis and carbon utilization related terms (GO:0009768; GO:0015976) (Fig. S2B).

### Transcriptional changes in CCM-related genes across cell types

We annotated 629 trinity unigenes with a known or hypothesized function in CCMs, including starch metabolism, circadian rhythm, and associated transcription factors as in (Christin *et al*., 2014; Ferrari *et al*., 2020; Gilman *et al*., 2021), (online Table S3). For most core CCM enzymes, unigenes were annotated to eudicot gene lineages as in (Christin *et al*., 2013, 2014; Moreno-Villena *et al*., 2017; Ferrari *et al*., 2020; Gilman *et al*., 2021). When available, eudicot gene lineage nomenclature is indicated after the gene names followed by a dash. 480 unigenes were kept after filtering lowly abundant unigenes; of those, 170 (35.42%) were DE across cell types, with 83 (48.82%) more abundant in bundle sheath and 87 (51.18%) more abundant in mesophyll. We found 307 (63.05%) CCM-related unigenes were DE over time, with 230 (74.92%) more abundant at 7h and 77 (25.08%) more abundant at 23h (Additional table S12). Finally, 75 (14.79%) CCM-related unigenes were DE across water regimen, with 61 (81.33%) more abundant in well-watered plants, and 10 (13.33%) more abundant in droughted plants.

Differential expression analyses of C4-related genes were consistent with expectations of C_4_ mesophyll vs. bundle sheath expression. In well-watered and droughted plants, the initial C_4_ carbon fixation module that included a carbonic anhydrase (*BCA-2E3*), *PEPC-1E1a’*, and an aspartate aminotransferase (*ASPAT-3C1*), occurred in mesophyll cells (mesophyll vs bundle sheath; MvsB log2FC > 1.5; *P*adj < 0.01; Module 1, Fig. 3-S2, Table S3). We identified a second bundle sheath specific (MvsB log2FC = -1.31; *P*adj < 0.01) *ASPAT* homolog (*ASPAT-1E1*) as a candidate gene representing the entry point for C_4_ metabolites into the CO_2_ decarboxylation/assimilation module in the bundle sheath that includes malic enzyme (*NADME-2E*), and the Calvin cycle (MvsB log2FC > -1.4; *P*adj < 0.01; Module 2, Fig. 3). We uncovered two malic enzymes (MEs) with C4-like expression, e.g., biased towards the morning and restricted to the BS cells. In addition to the primary *P. oleracea* ME used for C4-acid decarboxylation (mitochondrial malic enzyme *NADME-2E*) (Voznesenskaya *et al*., 2017), we found a chloroplastic ME (*NADP-ME-1E1b*) that may suggest an accessory decarboxylation route (Fig. 4). However, previous western blot analyses showed a low abundance of NADP-ME compared to NAD-ME *(41)*, suggesting that NADP-ME activity may be post-transcriptionally regulated. The expression of genes involved in the regeneration of PEP were restricted to the mesophyll (MvsB log2FC > 1.4; *P*adj < 0.01; Module 3, Fig. 3-S2). Among CCM-related genes, photorespiration genes were mainly expressed in bundle sheath (Module 9, Fig. S2), with the exception of chloroplastic D-glycerate 3-kinase (GLYK) which has been previously reported to be mesophyll specific *(42)*.

**Fig. 3.**
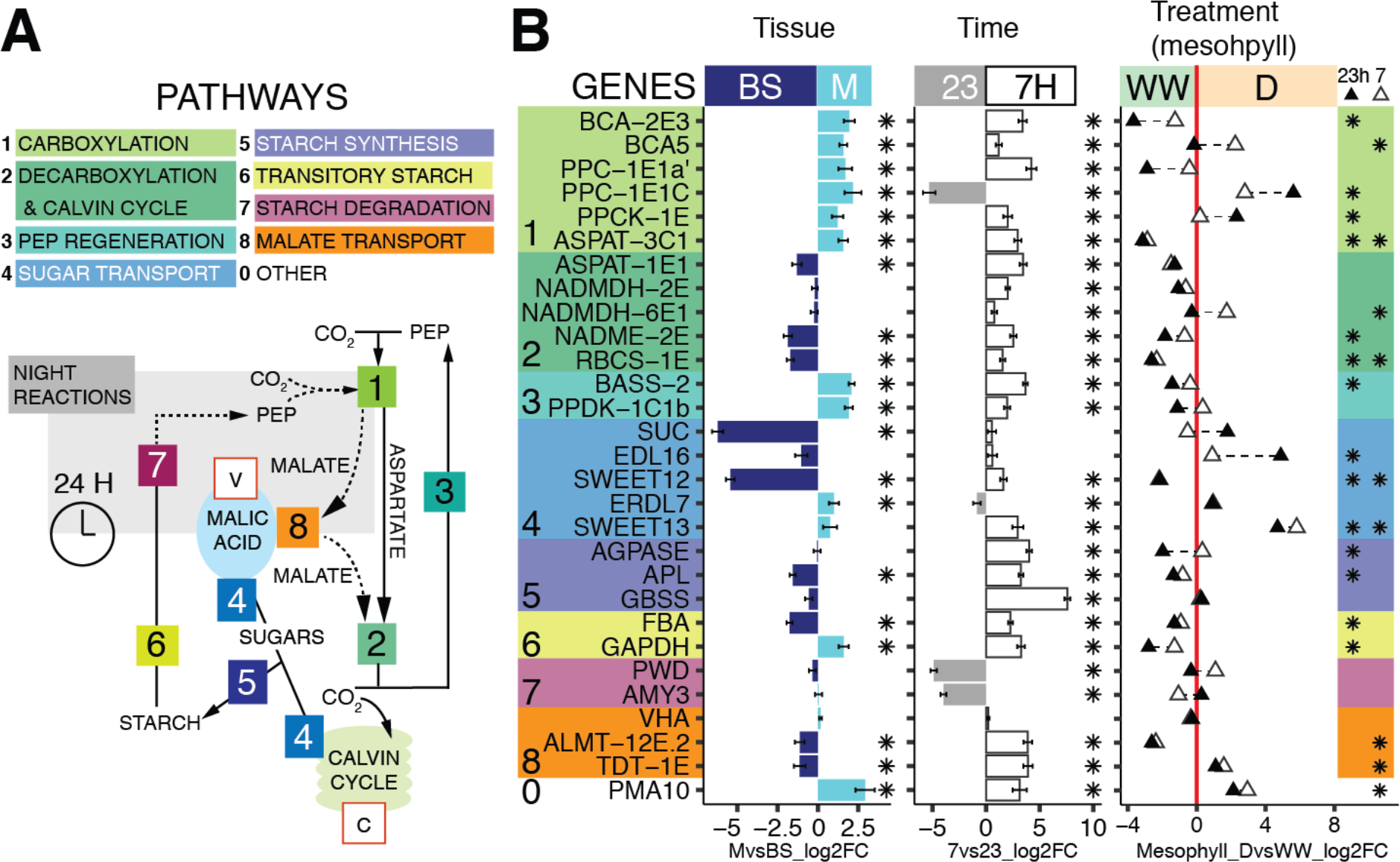
Differential gene expression of mesophyll and bundle sheath across experimental conditions. (A) Schematic of C4, CAM, and accessory biochemical pathways; solid and dotted lines indicate C_4_ and CAM routes of carbon concentration, respectively, the gray box contains night reactions, and key substrates are shown (Abbreviations: PEP, phospho*enol*pyruvate). Intracellular compartments are indicated within red boxes (Abbreviations: V, vacuole; C, chloroplast). (B) Differential transcript abundance (measured in log2 fold change, log2FC) of selected genes in mesophyll (M) relative to bundle sheath (BS) tissue (left panel) and at 07h relative to 23h (central panel) across LMD samples. Gene colour backgrounds reflect pathways in (A). In the third panel, white triangles indicate log2FC of 7h mesophyll samples in drought:D relative to 7h mesophyll samples well-watered:WW. Black triangles indicate log2FC of 23h mesophyll samples in drought compared to 23h mesophyll samples well-watered. A triangle in the WW region (negative log2FC, left to the red line) indicates higher expression during WW, while triangles within the D region (right to the red line) indicate higher expression in D. In all panels, asterisks indicate differential expression significance (*P*adj < 0.05).

**Fig. 4.**
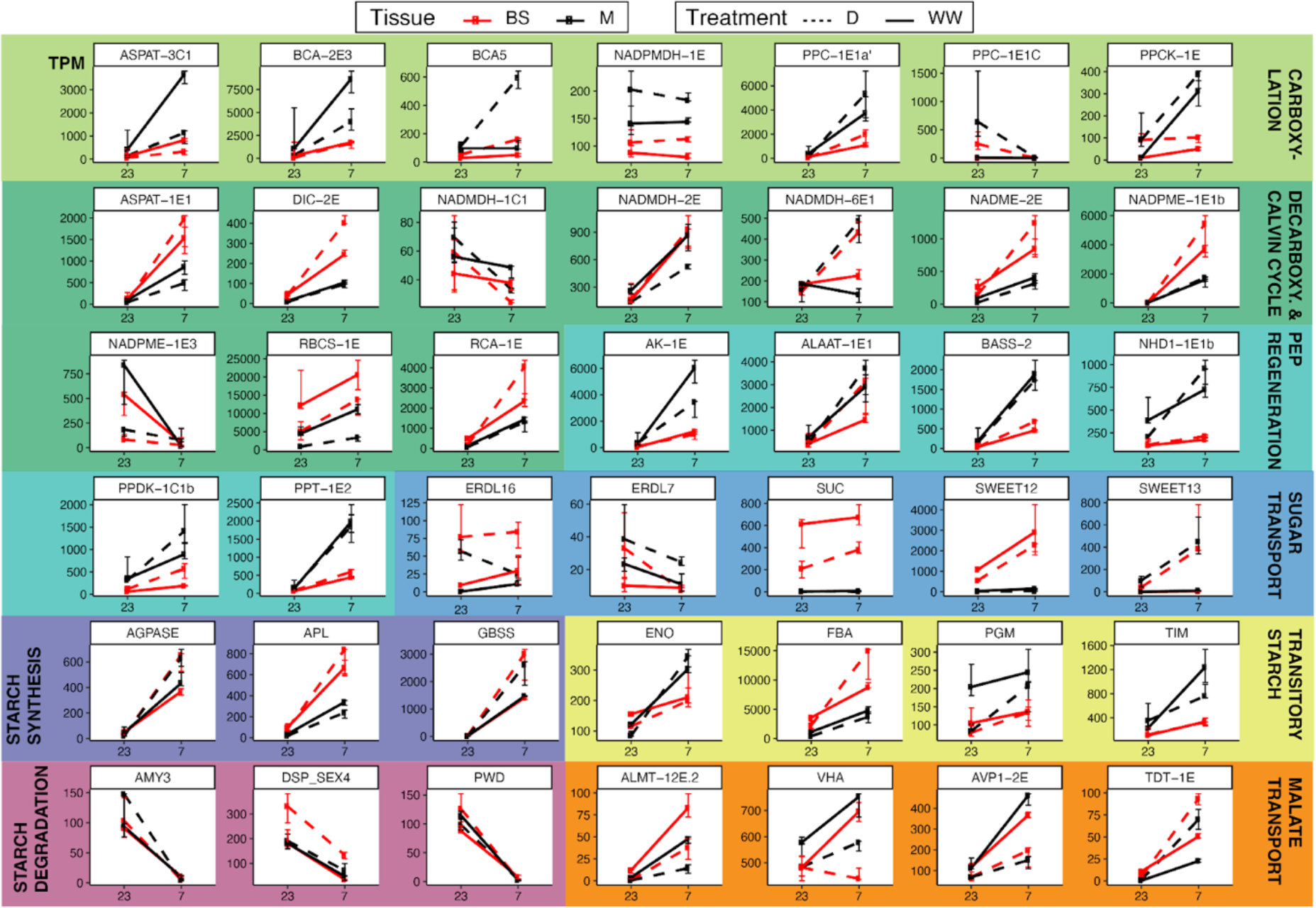
Transcription abundance of selected CCM-related genes. Median of transcripts per million (TPM, y-axis), across time points (7h and 23h, x-axis) in LMD mRNA libraries. Black and red lines indicate mesophyll or bundle sheath samples, respectively. Plain lines indicate watered and dotted indicate drought. Error bars represent the interquartile range of expression.

The ortholog identity and tissue localization of core C4-cycle genes generally remained the same in well-watered and drought-induced C_4_+CAM *Portulaca oleracea*. When CAM was induced, different homologs of carbonic-anhydrase (*BCA-2*), PEP carboxylase (*PEPC-1E1c*), and malate dehydrogenase (*NADMDH-6E1*) dramatically increased their mesophyll abundance (Mesophyll, drought vs well-watered; M_DvsWW log2FC > 1.7; *P*adj < 0.01; Fig. 3-4). In the mesophyll, the CAM-specific paralog *PEPC-1E1c* exhbited a 5.6 log2FC at night under drought, compared to well-watered day time samples, which had near-zero abundance (*P*adj < 0.01; Fig. 3-4). In parallel, the PEPC activating protein, PEPC kinase (*PPCK-1E*), increased in abundance at night in drought samples (M_DvsWW log2FC > 2.3; *P*adj < 0.01; Fig. 3-4).

Malate produced at night is purportedly actively pumped into the vacuole by an aluminum-activated malate transporter (*ALMT-12E2*) (Kovermann *et al*., 2007), which is driven by cation currents created by a vacuolar-type proton ATPase (*VHA*) (Smith *et al*., 1996) though neither showed increased expression during drought in any cell type (Module 8, Fig. S3). However, we found a sharp upregulation of the gene encoding the plasma membrane proton pump ATPase 10 (*PMA10*) in drought samples (M_DvsWW log2FC = 3; *P*adj < 0.05, Module 0, Fig. 3) restricted to the mesophyll (MvsBS log2FC = 2.9; *P*adj < 0.01; Fig. 3), but a role in CAM is only speculative. The mechanisms responsible for vacuolar malate efflux in CAM remain unknown (Borland *et al*., 2009), but the most plausible is the tonoplast dicarboxylate transporter (*TDT-1E*), whose expression increased during drought, particularly in mesophyll (M_DvsWW log2FC = 1.6; *P*adj < 0.01; Module 8, Fig. 3).

Unlike C_4_ plants, CAM plants generally produce PEP via degradation of starch and sugars; however, our sampling only showed significant up-regulation of the amyloplastic Phosphoglucan phosphatase *DSP4 (DSP_SEX4*) among starch and sugar degradation genes during CAM induction (Modules 6 and 7, Fig. 3-S3-S4). Temporally, starch degradation genes were more abundant at night (Module 7, Fig. 3-S2). Spatially, most of the transitory starch-related genes (Module 6, Fig. 3-S2) were more abundant in mesophyll during the day, with the exception of chloroplastic fructose-bisphosphate aldolase (*FBA* gene family, *ALFP* lineage; Table S3), which was restricted to bundle sheath (MvsBS log2FC = 1.8; Padj < 0.01; Fig. 23). FBA degrades fructose 1,6-bisphosphate into glyceraldehyde 3-phosphate (G3P), before G3P is further degraded to PEP in a series of glycolytic reactions *(as illustrated in Ferrari et al. (2020) figure 7b)* whose encoding genes were mainly expressed in the mesophyll during the day (Module 6, Fig. 3-S2). Tonoplast and chloroplast sugar transporters showed the highest expression specificity to bundle sheath (Module 5, Fig. 3,S2), suggesting a role in the Calvin cycle of C_4_ and C_4_+CAM plants. Other sugar transporters such as *ERD6-like (EDL16)* genes and the mesophyll restricted SWEET13 also increased in abundance with CAM induction (M_DvsWW log2FC > 5,8; Padj < 0.01; Fig. 2-S3). Our data suggests an increase in sugar transport linked to CAM induction, with G3P originating in the bundle sheath but the final steps of PEP regeneration mainly occurring in mesophyll.

We also found up-regulated genes related to cell wall architecture at night in CAM-induced plants: *EXPA2* and *SAU32* in the mesophyll and *NAP2* in bundle sheath (Table S4; Fig. S4). Together with the mesophyll-specific up-regulation of the ion transporters probable magnesium transporter (*NIPA7*) and plasma membrane-type ATPase 10 (*PMA10*), and bundle sheath-specific Mitochondrial uncoupling protein 4 (*PUMP4*) (Table S4; Fig. S4), we speculate that they may play a role modulating transport of starch and sugar across cells during CAM (Okumura *et al*., 2016). A gene involved in vacuolar processes (*VPE1*) in bundle sheath is also upregulated at night during CAM induction. The nocturnal activity of CAM is regulated largely by transcription factors (TFs) tied to the circadian clock. We detected expression shifts in TFs possibly linked to CAM induction and water stress responses, including increased diurnal expression of Transcription repressor *MYB4* in mesophyll and Phenylacetaldehyde reductase *PAR1* in bundle sheath, and decreased diurnal abundance of Transcription factor *BH062* in both cell types. During the night, we observed up-regulation of the TFs Scarecrow-like protein 15 (*SCL15*), Protein nuclear fusion defective 4 (*NFD4*), Homeobox-leucine zipper protein ATHB7, and probable *WRKY* transcription factor 7 (*WRKY7*) in both cell types, and Protein light-dependent short hypocotyls 3 (*LSH3*) in the mesophyll specifically (Table S4; Fig. S4).

### Visium spatial gene expression

To confirm LMD results, we captured and spatially tagged mRNA directly on leaf paradermal sections using the 10x Genomics Visium spatial transcriptomics platform (10X Genomics; Pleasanton, CA). *K*-means clustering of total gene expression grouped areas corresponding to mesophyll, bundle sheath, and water storage tissues (Fig. 3A). Genes with transcription estimates of less than 200 TPM in the LMD analysis were poorly represented in the Visium libraries. Transcript abundance mapped onto leaf-section micro-photographs revealed CCM and Calvin cycle genes were lowly abundant or undetected in water storage tissue, including both C4- and CAM-specific *PEPC* paralogs (Fig. 5-S5). Our Visium analyses confirmed the near absence of CAM *PEPC-1E1c* expression in well-watered samples and the day-night alternation of abundance of C_4_ and CAM *PEPC* paralogs in drought samples.

**Fig. 5.**
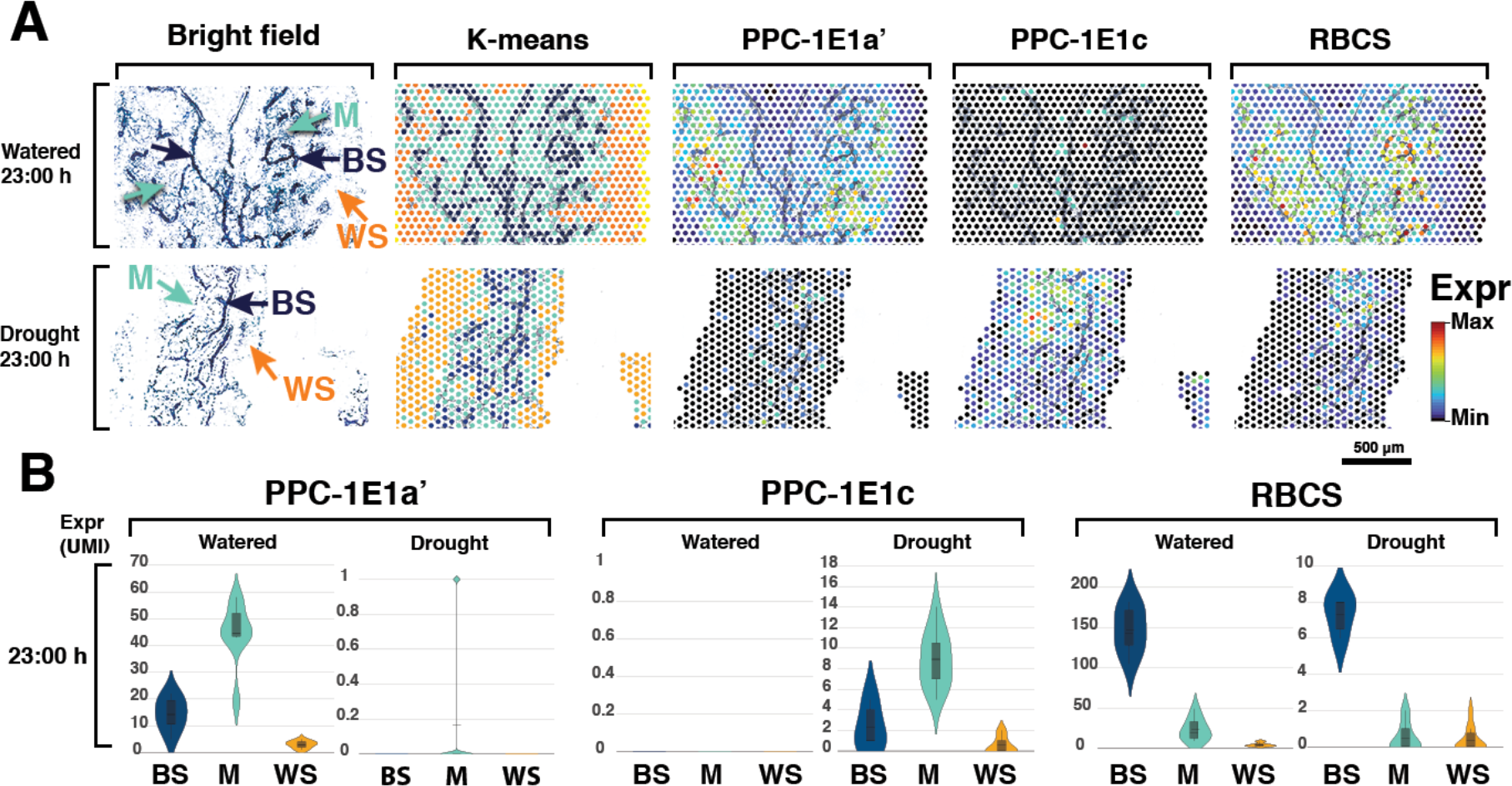
Visium spatial gene expression. (a) Microphotograph of a leaf paradermal section, *K*-means clustering of total gene expression, and abundance of the main C4, CAM and Calvin cycle carboxylases using the 10x Genomics Visium platform. *K-*means clustering of sampling spots corresponds to bundle sheath (BS, dark blue), mesophyll (M, light blue), and water storage (WS, orange) tissues; abundances of *PPC-1E1a’*, *PPC-1E1c*, and *RBCS* are shown relative to their observed unique molecule index (UMI) ranges. (b) Violin plots of transcript abundance in UMI across sample spots classified by tissue type in 23:00 h samples.

*PEPC* was mostly restricted to the mesophyll, with decarboxylation genes and Rubisco confined to the bundle sheath (Fig. 5-S5).

### Flux balance model

To further explore how C_4_ and CAM might be integrated, we built a two-cell two-phase model adapted from a highly curated plant core metabolic model (Shameer *et al*., 2018). The core metabolic model includes all major metabolic enzymes and reactions found to be highly conserved across a wide array of plant genomes, and represents a ‘core’ stoichiometric model, which captures the central carbon metabolism in leaves. Our new model represents how C_4_ and CAM could be integrated in a parsimonious manner from first principles of general plant metabolism. We used parsimonious flux balance analysis (pFBA) to model the efficiency of possible C_4_ and CAM configurations. In our models, metabolic fluxes were constrained by the rates at which metabolites were consumed or produced by each reaction and considered diel fluxes within mesophyll and bundle sheath cells as distinct compartments. All reactions were permitted to occur in both cells and at any time. The predicted fluxes for major reactions in the metabolic models were mostly consistent with the differential gene expression results (Fig. 6-S6), and most inconsistencies fell within the predicted variance of FVA (Fig. S7).

**Fig. 6.**
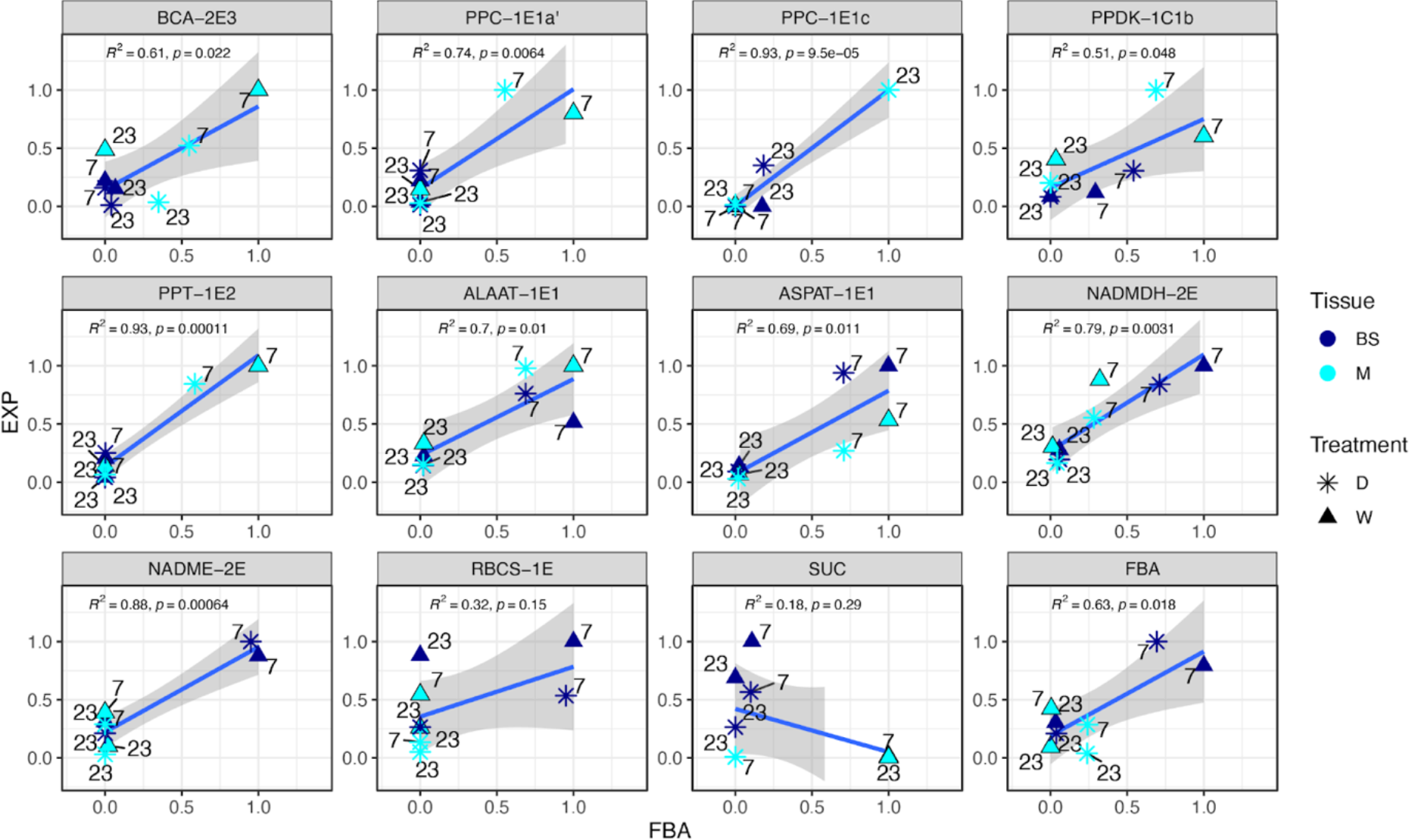
Correlations between predicted enzymatic fluxes and estimated gene expression in mesophyll and bundle sheath samples. Pearson correlation for individual enzymes between predicted reaction fluxes in the most efficient C_4_+CAM flux balance model (FBA) and the expression (EXP) of their encoding genes as the mean of transcript abundance in the LMD samples (transcript per million, TPM). Each data point corresponds to an experimental group of samples composed of drought (D) or wet conditions (W), mesophyll (M) or bundle sheath (BS) tissue and daytime (7h) or nighttime (23h). Time labels are shown next to each point. Expression and flux balance results are normalized by maximal values for each enzyme/gene. Regression lines are shown in blue and standard errors in grey.

Under well-watered conditions, the flux balance model predicted a C_4_ photosynthesis pathway with a small amount of CAM cycling in the bundle sheath; under drought conditions, the model predicted the emergence of a C_4_+CAM pathway (Fig. 6). While the model predicted a canonical C_4_ cycle, the predicted CAM cycle was a two-cell cycle spanning mesophyll and bundle sheath using both atmospheric and respired CO2, and a regular one-cell cycle in bundle sheath using respired CO2. For the two-cell CAM cycle, CO_2_ was assimilated by PEPC, converted to malate, and stored in the mesophyll vacuole. During the day, malate was transferred to and decarboxylated in the bundle sheath, where the C3 cycle was completed. All the essential C4-and CAM-related reactions (PEP carboxylation, carbonic anhydrase activity, Rubisco carboxylation, CO_2_ uptake, NAD-ME activity, and PEP dikinase regulation) had little variation in the FVA, indicating they were robust processes (Fig. S7). Most of the processes related to metabolite turnover (i.e., malate, OAA, ALA, ASP, PYR) were quite flexible (Fig. S7). Although pFBA predicted a solution with the lowest sum of fluxes, these fluxes exhibited great variation without affecting the efficiency of the system.

We further looked at whether blocking intercellular malate transfer and malate storage would affect the efficiency of the C_4_+CAM system. Blocking malate transfer did not affect the efficiency of the system under both well-watered and drought conditions (Table S7), because malate, ASP, and ALA transfer between mesophyll and bundle sheath were interchangeable, and therefore blocking one metabolite induced conversion to an alternative metabolite for intercellular movement. Furthermore, malate transfer may have the smallest enzymatic and metabolic cost in the C_4_+CAM system. Blocking malate storage in mesophyll or bundle sheath alone did not affect the efficiency of the system; however, blocking malate storage in both mesophyll and bundle sheath greatly reduced efficiency and introduced instability. Malate storage is a requirement for C_4_+CAM metabolism, but it is not necessarily important whether the malate is stored in mesophyll or bundle sheath.

In summary, under well-watered conditions, pFBA predicted a C_4_ pathway with only a very small amount of CAM in the bundle sheath; but under drought, pFBA predicted a C_4_+CAM system, consistent with our gene expression results (Fig. 7-S6-S7). Under drought, integrating malate from CAM into the C_4_ cycle maximized phloem photosynthate output with minimal enzyme investment compared to models where C_4_ and CAM cycles ran independently of one another. In the integrated model, most nocturnal CO_2_ fixation occurred in the mesophyll, and malate was stored in vacuoles of both cells. During the day, fluxes of malate from mesophyll vacuoles moved to the bundle sheath to enter the C_4_ decarboxylation module.

**Fig. 7.**
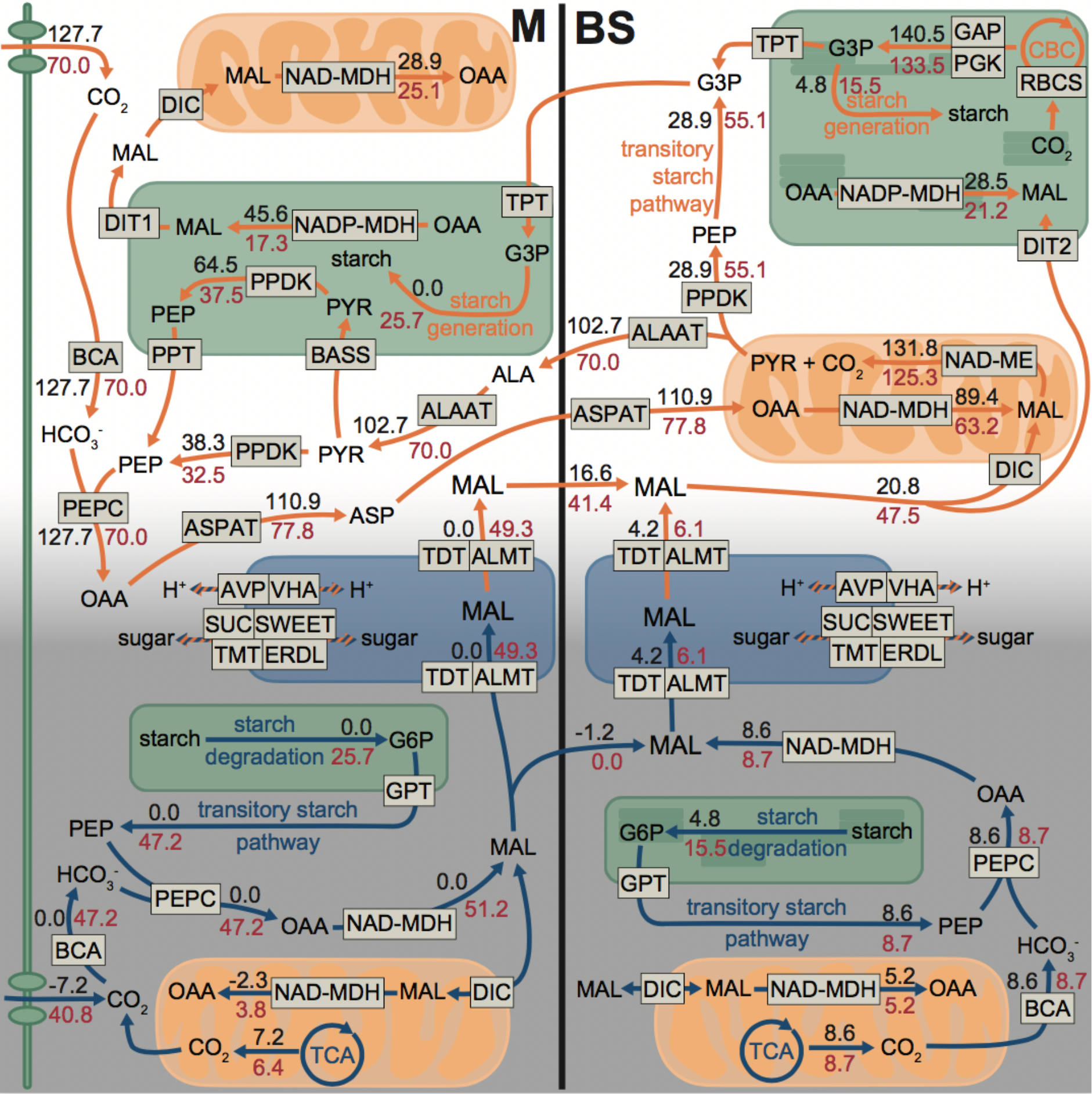
Biochemical flux results in the pFBA model of an integrated C_4_+CAM photosynthesis. Schematic of major metabolic fluxes related to C_4_ and CAM in the mesophyll (M) and bundle sheath (BS); orange and blue arrows indicate daytime and night-time reactions, respectively. Black numbers indicate fluxes under well-watered conditions (300 ppm internal CO_2_ concentration at daytime, no limit for night-time); red numbers indicate fluxes under drought conditions (70 ppm CO_2_ internal concentration at daytime. no limit for night-time). Minor fluxes are not shown for simplicity and therefore not all shown input and output fluxes are equal.

## Discussion

*Portulaca oleracea* is a rare C_4_ species capable of expressing CAM photosynthesis as a stress response to water deficit. This metabolic flexibility allows *P. oleracea* to achieve high rates of photosynthesis when resources are plenty but also to tolerate extreme drought for a relatively long period while maintaining the core carbon metabolism, breaking a fundamental trade-off in plant physiology. Previous studies identified candidate enzymes linked to C_4_ and CAM, but how the two pathways were spatially organized within *Portulaca* leaves remained unknown. We compared well-watered and drought-induced CAM-expressing *P. oleracea* plants and utilized the most recent advances in spatial gene expression analysis to describe for the first time a novel biochemical pathway that integrates C_4_ and CAM cycles into a single metabolic system.

Laser captured mesophyll and bundle sheath tissues revealed CAM and C_4_ operating in the same cells, confirmed by the Visium spatial gene expression analyses.

Traditionally, the spatial description of gene expression across different tissues via in-situ hybridization is necessarily limited to a handful of genes. Here, we estimated the expression of tens of thousands of genes in mesophyll and bundle sheath cells using laser capture microdissection. Furthermore, using the Visium (10X Genomics) spatial gene expression workflow, we mapped and visualized the expression of C_4_ and CAM related genes over paradermal anatomical sections of leaves. Spatial gene expression analysis at this scale is still quite rare in plants (Giacomello *et al*., 2017) and holds exceptional promise for addressing a diverse set of problems in plant molecular biology and function. Based on K-means clustering of similar gene expression, our Visum spatial gene expression analysis predicted the areas covered by mesophyll, bundle sheath and water storage cells across samples. Gene expression maps overlapping microphotographs of leaf sections were congruent with the findings in the laser captured cell samples: genes related to C_4_ and CAM PEPC activity were more abundant in mesophyll, while decarboxylation and Calvin cycle genes were biased to bundle sheath. In areas where water storage cells were present, metabolic activity appeared very low, and almost no transcripts were captured for any CCM related genes. Future work using spatial proteomics and metabolomic profiling will be decisive to confirm these results.

### A new photosynthetic pathway: CAM metabolic fluxes are connected to the C_4_ pathway for decarboxylation

We also built a two-cell two-diel phase plant metabolic model adapted from a highly curated model (Shameer *et al*., 2018) and used flux balance analysis (FBA) to test for the most efficient biochemical pathways in which C_4_ and CAM could operate, and correlated the results with our transcript abundance estimations. Our new model represents the integration of C_4_ and CAM based on first principles of general plant metabolism. The model predicted that a C_4_+CAM system would perform with higher efficiency under drought than a pure C_4_ alone; yet when drought conditions are alleviated, a pure C_4_ system has a higher efficiency than C_4_+CAM. This is congruent with observed *P. oleracea* behavior. Second, FBA models predict the cellular compartmentalization of the major C_4_ and Calvin Cycle reactions in agreement with our expression data, with the caveat that transcripts abundance of RuBisCO, NAD-MDH and NADME are higher in mesophyll than the model predicted (though still mostly expressed in bundle sheath). These results could be partially explained by an incomplete segregation of enzymes, or by the imperfect nature of laser microdissection for mesophyll and bundle sheath isolation. Third, under drought conditions, the integration and processing of malate from CAM into the C_4_ cycle maximizes phloem output of photosynthates with the least enzyme cost compared to models without integration of cycles. Under this model, most of the nocturnal CO_2_ fixation occurs in mesophyll and a fraction in bundle sheath and is stored in vacuoles of both cells. During the day, fluxes of malate from mesophyll reach bundle sheath to enter the C_4_ decarboxylation module. All together, we present for the first time a two-cell CAM system operating in *P. oleracea*, where CO_2_ is fixed by the C_4_ and CAM machinery in the same mesophyll cells but at distinct times, with CAM fluxes of malate entering the C_4_ cycle during the day in bundle sheath cells to be decarboxylated, and finally assimilated into sugars by Rubisco and the Calvin cycle.

### Abundant gene copies, ancestral succulence, and unusual venation architecture enabled the evolution of C_4_+CAM in *Portulaca*

Our analyses confirm that *P. oleracea* possesses an integrated C_4_+CAM photosynthesis, where initial C_4_ and CAM carbon fixation occurs in the same mesophyll cells over a 24-hour period, with decarboxylation and final CO_2_ assimilation restricted to the bundle sheath. Only a subset of core enzymes recruited distinct homologs for C_4_ vs CAM expression. Thus, the regulatory constraints that were proposed to prevent this coordination appear easily surmountable; not all genes require a temporal or spatial shift in gene expression, and those that do simply utilize alternative homologs. So why is the co-occurrence of C_4_ and CAM so rare, having only been identified in *Portulaca*, and quite recently, *Trianthema* (Winter *et al*., 2021)? It is possible that many C_4_+CAM species exist and have not been identified, as most known C_4_ species have not been investigated for CAM activity. At the same time, it is helpful to look at what *Portulaca* and *Trianthema* have in common. Both are mildly succulent plants that are unusual C_4_ members of clades with predominantly C3+CAM or CAM metabolism. *Portulaca* was ancestrally a C3+CAM species that evolved C_4_ in parallel several times (Ogburn and Edwards, 2013), and we predict that *Trianthema* will be shown to have followed a similar evolutionary trajectory. *Portulaca* achieved the high bundle sheath: mesophyll ratio needed for efficient C_4_ photosynthesis by evolving a three-dimensional leaf venation system while still maintaining tissue succulence (Ocampo *et al*., 2013) - thus traversing the potential antagonism of C_4_ and CAM anatomical requirements. We predict that anatomy may place a larger evolutionary constraint on the emergence of a C_4_+CAM system than genetic or metabolic coordination and that we are more likely to find additional C_4_+CAM species in mildly succulent C_4_ clades (e.g. Kadereit, Ackerly and Pirie, 2012) or C_4_ clades closely related to CAM-evolving lineages (e.g. Yang *et al*., 2012).

### A C_4_+CAM metabolism and global food security

Facultative CAM cycles in C3 plants help to maintain a positive carbon balance during stress (typically drought) (Winter, 2019). In combination with C4, a facultative CAM cycle affords elevated drought tolerance in a plant that can also achieve exceptionally high rates of photosynthesis, essentially circumventing the fundamental productivity-tolerance trade-off that constrains plant function (Grime, 1977). Future work should confirm the benefits of facultative CAM in *Portulaca* experimentally via gene editing. *Portulaca* is highly amenable to development as a model system for photosynthesis research (Callegari Ferrari *et al*., 2021): it has a short life cycle, is self-compatible, and a high-quality genome of *P. amilis* is already available (Gilman *et al*., 2021).

An integrated C_4_+CAM photosynthesis inspires future avenues for crop improvement and food security, as there has been an ongoing global initiative to engineer C_4_ or CAM pathways into C3 crops (Yang *et al*., 2015; Ermakova *et al*., 2020). Predictions of increasing evapotranspiration with warming climate point to the urgency of developing drought tolerant crops (DeLucia *et al*., 2019); but in maize, decades of selection for more robust crops resulted in significantly higher drought sensitivity (B. *et al*., 2014) Because maize, as a C_4_ plant, already possesses the ability to process malate for sugar production during the day, engineering a facultative CAM cycle may require only a handful of the changes that a full C3 to CAM or C3 to C_4_ transition would require. A C_4_+CAM maize could lead to lower-input, higher yield agriculture with a smaller carbon footprint at a global scale.

## Acknowledgments

We are grateful to Dr Emilie Guillon, at Yale University, for sharing with us her expertise with the microscopes. We also thank the Horsley and Wolenski Labs, at the Department of Molecular, Cellular, and Developmental Biology, Yale University, for sharing with us their laboratory instruments, and to Christopher Bolick at Yale’s Marsh Botanic Gardens, for the support with the experiment. Finally, we thank R. Shawn Abrahams, Anri Chomentowska, Kirstin Dion, Nora Heaphy, Joshua Randall, and Oluwatobi Oso for comments on earlier drafts that have improved this manuscript. This work was funded by the National Science Foundation (IOS-1754662 award to E.J.E). This work is supported by the NOAA Climate and Global Change Postdoctoral Fellowship Program (award #NA18NWS4620043B to H.Z)

## Author contributions

Conceptualization: JJMV, HZ, EJE

Methodology: JJMV, HZ, ISG, SLT, CYMC, EJE

Investigation: JJMV, HZ, SLT, CYMC

Visualization: JJMV, HZ, ISG, EJE

Funding acquisition: EJE, HZ

Writing – original draft: JJMV, HZ, EJE

Writing – review & editing: JJMV, HZ, ISG, SLT, CYMC, EJE

## Supplemental figure legends

**Fig. S1. Gene Ontology enrichment across bundle sheath and mesophyll and across watering regimens (extension of fig. 2).** 50 most significant Gene Ontology terms enriched across differentially expressed genes between mesophyll and bundle sheath. Barplots indicate the percentage of genes up-regulated of each GO term across (A) mesophyll and bundle sheath, (B) well-watered and drought, (C) 23h and 7h.

**Figure S2. Differential transcript abundance across cell types and experimental conditoins (extended fig. 3)** Differential transcript abundances (measured in log2 fold change, log2FC) of selected genes in mesophyll (M) relative to bundle sheath (BS) tissue (left panel), 07h relative to 23h (middle panel) and drought relative to well-watered (right panel) across LMD samples. Gene colour backgrounds correspond with pathways in the boxes on the top. In all panels, asterisks indicate significant differential expression (*P*adj < 0.05).

Figure S3. Effect of watering regimen in each cell type across time points (extension fig. 3).

Differential transcript abundance in droughted:D relative to well-watered plants:WW (measured in log2 fold change, log2FC) of selected genes. Gene colour backgrounds correspond to pathways in the boxes on the top. Triangles on the left panel represent relative abundance of D mesophyll samples vs WW mesophyll, in 7h samples (white triangles) and in 23h samples (black triangles). Right panel shows the same abundance comparisons using only bundle sheath samples. A triangle in the WW region (negative log2FC, left to the red lines) indicates higher expression during WW, while triangles within the D region (right to the red line) indicate higher expression in D. Asterisks indicate significant differential expression (*P*adj < 0.05)

**Fig. S4. Transcription abundance of CCM-related genes and genes with no known role in C_4_ or CAM (extension of Fig. 4).** A) Transcription abundance of CCM-related genes listed in table S3. B) Genes with no known role in C_4_ or CAM listed in table S4. Median of transcripts per million (y-axis), across time points (x-axis) in LMD mRNA libraries. Black and red lines indicate mesophyll or bundle sheath, respectively. Plain lines indicate watered and dotted indicate drought. Error bars show the interquartile range of expression.

**Fig. S5. Visium spatial gene expression (extended fig. 5)**. The first row shows frozen leaf specimens at the moment of cryo-sectioning. The second row shows microphotographs of leaf paradermal sections under bright field (left) and K-means clustering of total gene expression (right). Successive rows show abundance of the main CCM-related genes using the 10x Genomics Visium platform. K-means clustering of sampling spots corresponds to bundle sheath (BS, dark blue), mesophyll (M, light blue), and water storage (WS, orange) tissues; abundances are shown relative to their observed unique molecule index (UMI) ranges.

**Fig. S6. Correlations between predicted enzymatic fluxes and estimated gene expression in mesophyll and bundle sheath samples** (related to fig. 6) Pearson correlation between z-score normalized (mean set to 0, SD set to 1) pFBA results and transcript abundance. The transcript abundances from different orthologs used in the same biochemical reactions were added together to be compared with the pFBA results. 23_BS_D: night-time flux in bundle sheath under drought; 23_BS_W: night-time flux in bundle sheath under well-water; 23_M_D: night-time flux in mesophyll under drought; 23_M_W: night-time flux in mesophyll under well-water; 7_BS_D: daytime flux in bundle sheath under drought; 7_BS_W: daytime flux in bundle sheath under well-water; 7_M_D: daytime flux in mesophyll under drought; 7_M_W: daytime flux in mesophyll under well-water;

**Fig. S7. Flux variability analysis (FVA) of mesophyll and bundle sheath with diel variation and under drought and well-water conditions (related to Fig. 7).** Daytime: orange arrows and white background; night-time: blue arrows and grey background.

Reactions that occur during the day-night transition are shown with striped arrows. M: mesophyll; BS: bundle sheath. Black numbers: well-water condition (300 ppm CO_2_ concentration at daytime and and no limit for night-time); red number: drought condition (70 ppm CO_2_ concentration at daytime and no limit for night-time).

Table S1 (related fig. 1A).

Leaf titratable acidity analysis results. Microequivalents H+ (μEq H+) per gram fresh mass was calculated as volume titrant (μL) ⨉ titrant molarity (M) / tissue mass (g).

Table S2 (related to fig. 2).

LMD-RNA sequencing and read mapping statistics. BS: bundle-sheath cells, M: mesophyll cells, n_: number of reads. T: time; Ind: Plant individual; n_raw: number of raw reads; n_filtered: number of reads after filtering; n_aligned: number of aligned read to the transcriptome; %_al: percentage of reads aligned to the transcriptome.

**Table S3. (related with figs. 3-4-5-S2-S3).** Annotation of selected genes with a role or potential role in CCM.

**Table S4. (related with fig S4**)Annotation of selected genes differentially expressed across experimental variables without a known role in CCM.

**Table S5. (related with visium spatial gene expression results)** Sequencing and reads mapping statistics across Visium mRNA libraries.

**Table S6. (related with figures 6-7)**. Sensitivity analysis of pFBA results of major reactions for Rubisco carboxylation to oxygenation ratio (*V*c/*V*o) for C_4_ in bundle sheath.

**Table S7.(related with figures 6-7)** pFBA results for major reactions of additional modelling scenarios in mesophyll (M) or bundle sheath (BS) at daytime (day) and night time (night). Major reactions are Phloem output, PEP carboxylation (PEP), Rubisco carboxylation (Rubisco) and CO_2_ absorption (CO2). All the scenarios are modelling under the drought condition. Scenario 1): blocking malate transfer between mesophyll and bundle sheath (bMalT). Scenario 2):blocking malate storage in mesophyll (bMS), bundle sheath (bBSS), or both (bMBSS). Scenario 3): CAM with C3 or C4anatomy. C3+CAM: both C3and CAM activity (both daytime and night time CO_2_ uptake) allowed with a C3 anatomy (CO2directly diffuses into mesophyll, bundle sheath is considered an inner mesophyll with a longer distance to stomata); CAM_C3: only CAM (night-time CO_2_ uptake) occurs with C3anatomy ; CAM_C4: CAM process (night-time CO_2_ uptake) with C4anatomy (CO2 cannot directly diffuse into bundle sheath); C_4_+CAM_C4: C_4_ and CAM (both daytime and night time CO_2_ uptake) can occur with C4anatomy (CO2 can not directly diffuse into bundle sheath), which can be used as the reference for all the above scenarios.

## Methods

### RESOURCE AVAILABILITY

#### Lead contact

Further information and requests for resources should be directed to and will be fulfilled by the lead contacts, Jose J. Moreno-Villena, Haoran Zhou

#### Materials availability

This study did not generate new materials

#### Data and code availability

- Table S8 to S11 (.xlsx) available on Dryad DOI https://doi.org/10.5061/dryad.931zcrjm6
- Visium loupe files Data S1 to S6 (.loupe) available on Dryad DOI https://doi.org/10.5061/dryad.931zcrjm6 Temporary access to Dyrad https://datadryad.org/stash/share/WnJbocSWXZePuSSq9AIhE1LG4-8nxDrB1FuDVAhEZsE
- All short sequencing reads are available in the GeneBank Sequence Read Archive (SRA) BioProject: PRJNA774250
- The scripts used for the data analyses are available on https://github.com/josemovi/Portulaca-spatial-C4-CAM

Any additional information required to reanalyze the data reported in this paper is available from the lead contact upon request

### EXPERIMENTAL MODEL AND SUBJECT DETAILS

*Portulaca oleracea* plants were grown from seeds in a greenhouse (voucher accession PO-russ2018) in soil mix FAFARD GROWING MIX #2 3.8CF, composed of 70% peat moss, 20% perlite, 10% vermiculite, starter nutrients, limestone, and a wetting agent. https://www.parisfarmersunion.com/product-p/0236930.htm

### METHOD DETAILS

#### Methods I: Spatial Gene Expression

##### Plant material, drought experiment and sampling

Two *Portulaca oleracea* plants were grown at the greenhouse facilities at the Marsh Botanical Garden, Yale University, New Haven, CT, USA. During the experiment, temperature was maintained at an average of 27°C and 22°C during the 15 hours of light and 8 hours of dark a day, respectively. Natural light intensity at plant level peaked around 1600 mol · m-2 · s-1 (photosynthetically active radiation, PAR) at the center of the light period.

The light period began at ∼5h (ending ∼20h), and leaf samples were taken at 7h, 19h, and 23h under well-watered (WW) conditions, and again after 7 days of complete water withholding (drought, D). Leaves used for titratable acidity assays were placed in 1mL Eppendorf tubes and flash frozen in liquid nitrogen; for cryosectioning, another 3-4 whole leaves were immersed in optimal cutting temperature compound (OCT) in plastic histology molds, and rapidly placed on an aluminium block partially submerged in liquid nitrogen to flash freeze. All samples were stored at -80 °C.

To assess nocturnal accumulation of malate, leaves from 7h and 19h samples from both WW and D plants were boiled in 60mL of 20% EtOH. After half of the volume evaporated, distilled water was added to return the initial 60mL; this process was repeated twice. The final 30 mL solutions were cooled to room temperature and titrated to a pH of 7.0 using 0.002M NaOH. Acidity, measured as μEq H+ per gram fresh mass, was calculated as volume titrant (μL) ⨉ titrant molarity (M) / fresh tissue mass (g).

##### Cell-specific laser microdissection, RNA isolation and sequencing

Laser microdissection (LMD) relies on cell identification for cell-specific capture. Here, we targeted mesophyll cells (M) and bundle sheath (BS) cells for RNA sequencing. We chose to use fresh frozen tissue, as opposed to fixed tissue and/or embedded in paraffin, to avoid potential RNA degradation. Although freezing can distort the leaf anatomy, bundle sheath cells are easily identified by their dense chloroplasts and proximity to veins, and mesophyll cells by sparse chloroplasts and calcium oxalate crystals (Fig. 1-D).

Frozen OCT moulds containing fresh leaves were moved from -80 °C to -20 °C inside a cryomicrotome (CM 3050S, Leica Biosystems; MA, US) to acclimate for 30 min. We obtained paradermal sections by cutting leaves parallel to the longitudinal axis with the cryomicrotome. Paradermal sectioning results in larger areas of BS and M tissues compared to cross-sectioning. Sections 12 µM across were laid on PEN membrane glass slides for UV laser microdissection (ThermoFisher Scientific, MA, US) and stored at -80 °C. At the time of laser microdissection, slides were brought to room temperature inside a desiccant box containing silica gel for ∼10 minutes. Room temperature dry slides were laser cut using a Laser MicroDissection scope (LMD7000, Leica, Leica Biosystems; MA, US) with laser parameters: Power 50, Aperture 9, Speed 4, and 20X microscope magnification. This relatively wide cutting laser ensured clear separation of BS and M tissues. More than 10 areas containing tens of cells each, were captured and accumulated in each sample of each tissue type. M or BS samples were captured into separate caps of 0.5 mL micro-tubes filled with 20 uL of extraction buffer (Arcturus PicoPure RNA isolation kit, ThermoFisher Scientific; MA, US).

RNA was isolated using PicoPure kits (Arcturus PicoPure RNA isolation kit, ThermoFisher Scientific; MA, US), with additional on-column DNAse digestion (RNase-free DNase, Qiagen; Germany). Extraction quality was evaluated using a Tapestation DNF-472T33 (Agilent Technologies; CA, US) with the high sensibility assay HS Total RNA, analysis mode: Plant RNA. Only extractions with RNA profiles clearly exhibiting the two plant-cell ribosomal RNA peaks and the two chloroplasts rRNA peaks were sequenced (Fig. 1S-A). We used SMART-Seq v4 Ultra Low Input RNA Kits for sequencing (Takara Bio Group; Japan), which first generated high-quality cDNA from ultra-low amounts of total RNA and then high-quality Illumina sequencing-ready libraries. The SMARTer anchor sequence and poly-A sequence served as universal priming sites for end-to-end cDNA amplification of mRNA. We used 250pg of total RNA input with 14 PCR cycles for cDNA synthesis and 10 PCR cycles for library amplification, following the manufacturer’s guidance. mRNA libraries were pooled and sequenced on a Novaseq6000 system (Illumina; CA, US) to generate ∼25 million 100bp paired-end reads per library. Library preparation and sequencing were done at the Yale Center for Genome Analysis, New Haven CT, USA.

##### Spatial whole transcriptome sequencing

To obtain near-cellular resolution of gene expression across entire leaf paradermal sections, we used the Visium Spatial Gene Expression platform (10X Genomics; CA, US) following the manufacturer’s directions. The Visium platform captures mRNA released from a tissue section fixed to a slide, spatially tagging each mRNA molecule prior to sequencing to later map their position and abundance over a corresponding microphotograph of the tissue section. We followed the Tissue Optimization workflow to confirm the compatibility of our leaf tissues with the solution and to optimize the permeabilization conditions to release the maximum mRNA in the shortest time. First, using a cryomicrotome, 12 µM paradermal sections were obtained from flash frozen leaves and each placed within the frames of one of the eight, 6.5 x 6.5 mm capture areas on an optimization slide. Sections were fixed with methanol and stained with hematoxylin and eosin (H&E). The slide was scanned under a bright field using a microscope (Axio Imager.M1; Zeiss, Germany) with a magnification of 10X per tile for comparison with final fluorescent images. We used permeabilization test times of 3, 6, 12, 18, 24, and 30 minutes in each capture area, including negative (tissue section not exposed to permeabilization reagents) and positive controls (stock isolated plant mRNA). cDNA was generated from the mRNA that bound to probes on the slide during permeabilization, using fluorescently labelled nucleotides. Tissue was then enzymatically removed from the slide, and the remaining fluorescently labelled cDNA was analyzed to select the optimal permeabilization time based on the fluorescence of each test area. Using the fluorescence capacity of the microscope (TRITC filter cube - filter Rhod - 800 ms exposure time), we observed maximum fluorescence in the corresponding sample area after 12 minutes of permeabilization. The software ZEN 2.6 (Zeiss; Germany) was used to control the microscope and stitch the microphotograph tiles.

Once the tissue permeabilization conditions were optimized, a total of eight sections, comprising two technical replicates at 7h and 23h under well-watered and drought conditions, were used for spatial mRNA sequencing. Each replicate was placed on one of two slides containing 4 capture areas. Each area contained 5,000 barcoded capture spots that were 55 μm in diameter - 100 μm centre to centre between spots. Spots were populated with primers that included Illumina sequencing primers, unique spatial barcodes per spot, unique molecular identifiers (UMI), and 30 nt poly(dT) sequences to capture poly-A mRNA. Tissue sections were processed as in the optimization steps: cryosectioning, followed by fixation, staining, slide imaging, and permeabilization. The mRNA released from overlying cells was captured by the primers on the spots and incubated with a mix containing reverse transcription reagents, producing spatially barcoded cDNA. After the second strand was synthesized, a denaturation step released it from the slide, and cDNA was transferred from each capture area to a corresponding tube for amplification and library construction following manufacturers specifications. Amplification, library preparation, and sequencing was done at the Yale Center for Genome Analysis, following the manufacturer’s directions, with a sequencing depth of at least 50,000 read pairs per spot in the capture area.

We mapped reads from both LCM and Visium experiments to a *de novo Portulaca oleracea* transcriptome assembly from (Gilman *et al*., 2021), Before that, we reduced the dataset to contigs containing coding sequences. For that, we first used TransDecoder v5.5.0 [http://transdecoder.github.io] to predict coding sequences within the transcripts, providing the software with precomputed BLASTX (Camacho *et al*., 2009) alignments to the UniProt protein sequences database (Consortium, 2019) to improve prediction. Next, we used CD-HIT v4.8.1 (Li and Godzik, 2006) to cluster coding sequences sharing 95% identity, retaining one representative. The indexes of the remaining sequences were used to subset the initial transcriptome assembly, yielding a reduced transcriptome for read mapping. Transcripts were functionally annotated using Trinotate v3.2.1 [http://trinotate.github.io] selecting the best Blast hit (lowest e-value score) of all the possible alignments. Annotation of transcripts into gene lineages for the gene families with a role in carbon concentrating mechanisms (CCMs) and starch metabolism was perfomed in (Gilman *et al*., 2021) as with gene lineage notations as in (Christin *et al*., 2013, 2014; Moreno-Villena *et al*., 2017; Ferrari *et al*., 2020; Gilman *et al*., 2021).

mRNA reads from mesophyll and bundle sheath tissues captured using laser microdissection were filtered and trimmed using Trimmomatic-0.39 (Bolger, Lohse and Usadel, 2014): 10 bp were removed from the beginning of the read; Illumina adapters, poly-A tails, and SMART-Seq primers were removed from the reads; a 5 bp sliding window trimming approach was used to clip the read at the 5’ end where the average quality score within the window fell below 20; single, low-quality bases from the beginnings and ends were also clipped. Only reads longer than 18 bp were kept after filtering. To quantify abundance, reads were mapped to the transcriptome assembly using the pseudo-aligner Kallisto v-0.45.0 (Bray *et al*., 2016) with 100 bootstrap replicates.

We used DEseq2 (Love, Huber and Anders, 2014) to test for gene differential expression (DE) across experimental variables. To generate Trinity unigene-level counts (as opposed to transcript-level counts), we used Sleuth (Pimentel *et al*., 2017) in ‘gene_mode’ to compile the Kallisto transcript counts into unigene counts. The resulting matrix of estimated counts was used to measure DE with DEseq2 in R (R Core Team, 2018). To estimate the magnitude and direction of the change in expression (measured as log2 fold change) and to statistically test for DE unigenes across cell types, time points, and water status, we included those variables and controlled for sequencing batch and plant individuals using all samples for DEseq2 normalization. Additionally, to specifically measure the effect of drought on each cell type at each time point, we separated our counts matrix by 7h and 23h and ran the DEseq2 function for each partition, including in these cases a cell type and water status interaction term for normalization. The P-values obtained from each test were adjusted (P-adjusted) for multiple testing by means of false discovery rate using the Benjamini and Hochberg method (Benjamini and Hochberg, 1995). We then tested for enrichment of gene ontologies (GO) terms in the lists of significant DE unigenes from each test (P-adj < 0.05).

First, we extracted the corresponding GO terms from the Trinotate analysis, and then used Fisher’s exact test, as implemented in TopGO (Alexa, Rahnenfuhrer and Lengauer, 2006), using the GO terms from all unigenes as the background ‘gene universe’.

##### Visium Spatial Gene Expression analysis

Reads obtained from tissue sections using the Visium Spatial Gene Expression platform were processed using the analysis pipelines implemented in Space Ranger V1.1.0 (10X Genomics) [https://support.10xgenomics.com/spatial-gene-expression/software/pipelines/latest/what-is-space-ranger]. We used the ‘spaceranger mkref’ to construct a reference transcriptome, parsing the exon unigene information generated by Transdecoder and the sequence of the longest isoform by unigene. We mapped reads to the most highly expressed unigene per *Portulaca* CCM-related gene lineage when possible, or per gene family, from the above LCM analysis, a total of 140 sequences. We manually aligned the bright field microphotograph based on fiducial markers and selected the areas where capture spots covered tissue. We then ran ‘spaceranger count’ to filter-trim reads and align them to the reference transcriptome, with default parameters. Unique molecular identifiers (UMI) were later used to correct and estimate counts of aligned reads per unigene per spatial capture spot. Principal components analysis (PCA) and *K*-means clustering of spots by expression similarity was performed automatically in Space Ranger and visualized in the Loupe Browser 5.0 (10X genomics).

#### Methods II: Flux balance model

To further explore how C_4_ and CAM might be integrated, we built a two-cell two-phase model adapted from a highly curated plant core metabolic model (Shameer *et al*., 2018). The core metabolic model includes all major metabolic enzymes and reactions found to be highly conserved across a wide array of plant genomes, and represents a ‘core’ stoichiometric model, which captures the central carbon metabolism in leaves. Our new model represents how C_4_ and CAM could be integrated in a parsimonious manner from first principles of general plant metabolism.

##### Model construction

Previous models only considered C3 or CAM metabolism in a single-cell model, or C_4_ metabolism in a two-cell model, but without diel variation (Shameer *et al*., 2018, 2020; Blätke and Bräutigam, 2019)). In order to consider the C_4_ and CAM cycles together, we built a new metabolic model including both mesophyll and bundle sheath cells and diel-flux changes. Our model is an extension of the (Shameer *et al*., 2018) model, which considered diel-flux and charge balance in each cellular compartment. The advantage of including charge balance is that it considers the effects of organellar pH on the metabolites’ charge states, which is important for the CAM acidification process. Starch was allowed to accumulate in the plastid, and sugars (glucose, sucrose, and fructose), carboxylic acids (malate, citrate, and isocitrate), proteinogenic amino acids, and nitrate were allowed to accumulate in the vacuole for storage and use during both day and night. We duplicated all the reactions to represent bundle sheath and mesophyll cell types individually. We then added transport reactions between bundle sheath and mesophyll for daytime and night-time conditions. We allowed particular metabolites to be transferred between bundle sheath and mesophyll based on previous studies (de Oliveira Dal’Molin *et al*., 2010; Mallmann *et al*., 2014; Arrivault *et al*., 2017; Shaw and Cheung, 2019) malate, aspartate, alanine, pyruvate, G3P, DHAP, PEP, GAP, 2-PG, sucrose, sulfate, ammonium, nitrate, and CO2. We also added charge- and proton- balances for these reactions following the method of (Shameer *et al*., 2018). Our model had the following four first-order compartments: daytime mesophyll, night-time mesophyll, daytime bundle sheath, and night-time bundle sheath. Under each of these first-order compartments, there were the following second-order compartments where corresponding biochemical reactions happen: plastid, peroxisome, vacuole, thylakoid, mitochondrion, extracellular, cytoplasm, mitochondrial intermembrane space, and inner membrane spaces. We also made the following revisions to the model: allowed proton and oxygen transfer between day and night, fixed Mannan biosynthesis reactions, and allowed protons to be transferred reversibly between cytoplasm and mitochondria, vacuole, extracellular space. Thus, our model contains all of the major biochemical reactions related to C3, C4, and CAM photosynthesis. Our model is available on GitHub

##### Modelling constraints

We used the COBRApy package (Ebrahim *et al*., 2013) to perform the flux balance analysis. The non-growth associated maintenance costs were controlled by constraining an ATPase and three NADPH oxidase pseudo-reactions (cytoplasmic, mitochondrial, and plastidic) to a 3:1 (ATP hydrolysis:NADPH oxidase) ratio, based on (Cheung *et al*., 2013) and used by previous studies (Shameer *et al*., 2018, 2020). The maintenance costs, which were represented by ATPase fluxes, had a linear relationship with photon uptake fluxes (Shameer *et al*., 2018). We used a photon uptake flux of 800 μmol m-2 s-1, similar to light levels in our greenhouse experiments described above, and a corresponding non-growth associated maintenance ATPase flux of 33.66 μmol m-2 s-1. We also set an anatomical constraint that bundle sheath cells occupied 30% of the leaf area and mesophyll cells 70%. All the fluxes in mesophyll and bundle sheath were adjusted according to the above ratios. Rubisco carboxylation to oxygenation ratio was set to 3:1 in mesophyll, but 10:1 in bundle sheath due to lower diffusion of O2 and the carbon concentration mechanism (Shaw and Cheung, 2019). We used a relatively lower Rubisco carboxylation to oxygenation ratio compared to other studies of C_4_ models, but sensitivity analysis indicated that our results were robust to varied and higher ratios (Table S6). Finally, as *P. oleracea* primarily uses NAD-type C_4_ (Gutierrez, Gracen and Edwards, 2004; Voznesenskaya et al., 2010) we constrained our model to use NAD-ME as the decarboxylation enzyme and accordingly blocked the transfer of pyruvate, PEP, 2PG, and GAP between mesophyll and bundle sheath (Furbank, 2011; Khoshravesh *et al*., 2020)We did not constrain any other major biochemical reactions related to C3, C4, or CAM.

##### Modeling scenarios

We parameterized the models with the growth environmental conditions in the greenhouse experiment, and performed the pFBA. The primary objective function maximized phloem output, which meant letting the model dictate which pathways were most efficient (i.e., generate the highest phloem output) under given environmental conditions. The secondary objective function minimized the absolute sum of fluxes, which was a proxy for reducing enzymatic costs while fulfilling the primary objective function (Töpfer *et al*., 2020).We then performed flux variability analysis (FVA) to examine the potential flux space without affecting the primary objective in pFBA analysis. We first let the model freely determine the potential pathways among C3, C4, and CAM with two objectives, without adding additional constraints, and under different environmental conditions. By mimicking stomatal conductance by setting different internal CO_2_ concentrations for the same model, we could predict whether and how C_4_ and CAM were integrated under well-watered (300 ppm CO_2_ concentration at daytime) and drought conditions (70 ppm CO_2_ concentration at daytime). We did not constrain night-time CO_2_ uptake and let the model freely determine it.

We tested several scenarios related to malate transfer and storage: 1) blocking malate transfer between bundle sheath and mesophyll; 2) blocking malate storage in mesophyll; 3) blocking malate storage in bundle sheath; and 4) blocking malate storage in both mesophyll and bundle sheath.

Lastly, we performed alternative modeling scenarios related to C_4_ anatomy. We revised the constraint of CO2-proof bundle sheath cells (typical C_4_ anatomy) by allowing CO_2_ to diffuse into bundle sheath, but maintained the same atmospheric CO_2_ as in modeling scenarios with CO2-impermeable bundle sheath. In order to set a CAM scenario, we set daytime CO_2_ at 0 and the Rubisco carboxylation to oxygenation ratio to 5.15, following (Shameer *et al*., 2018). All the other settings and constraints were the same for C3, C4, and CAM scenarios.

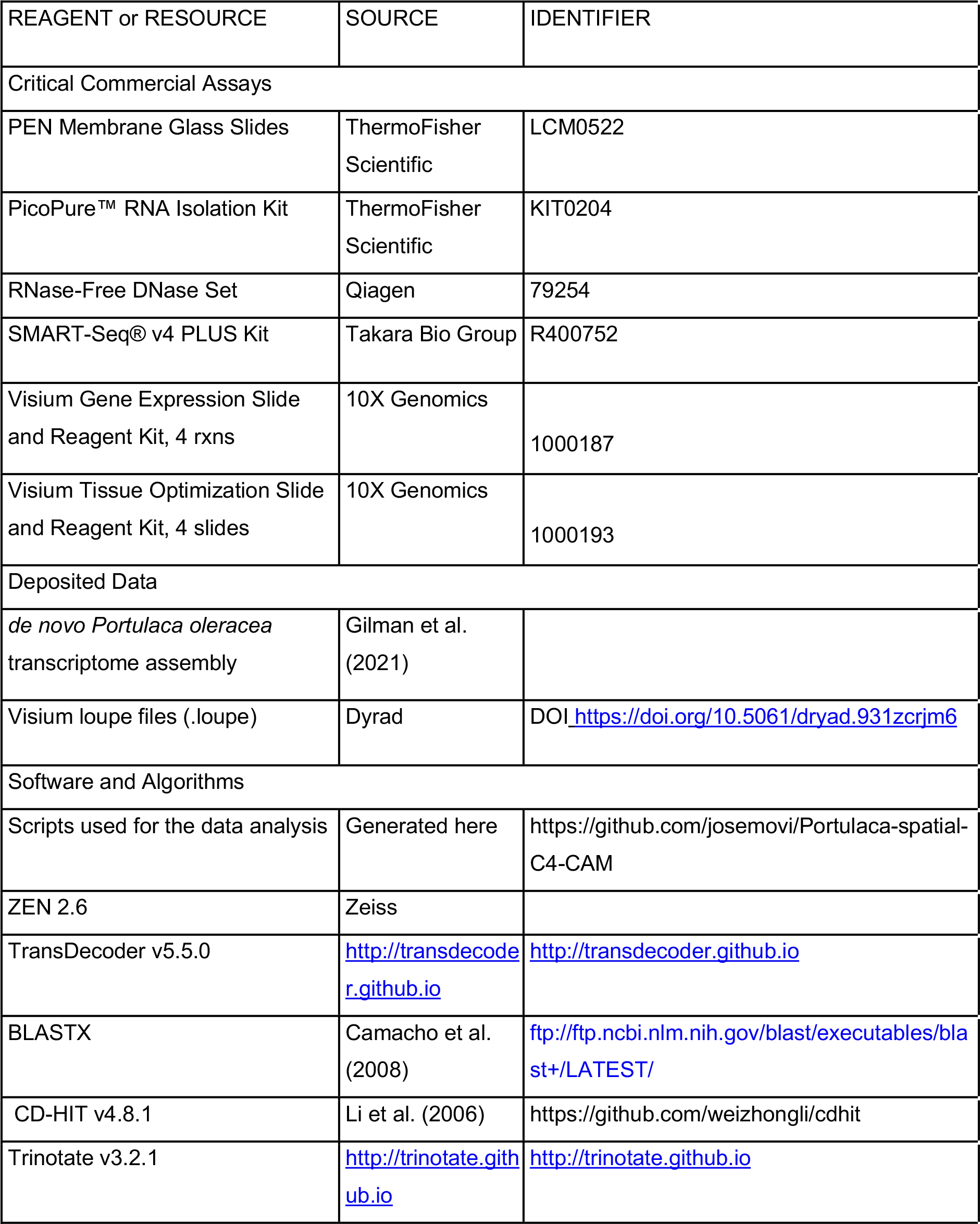
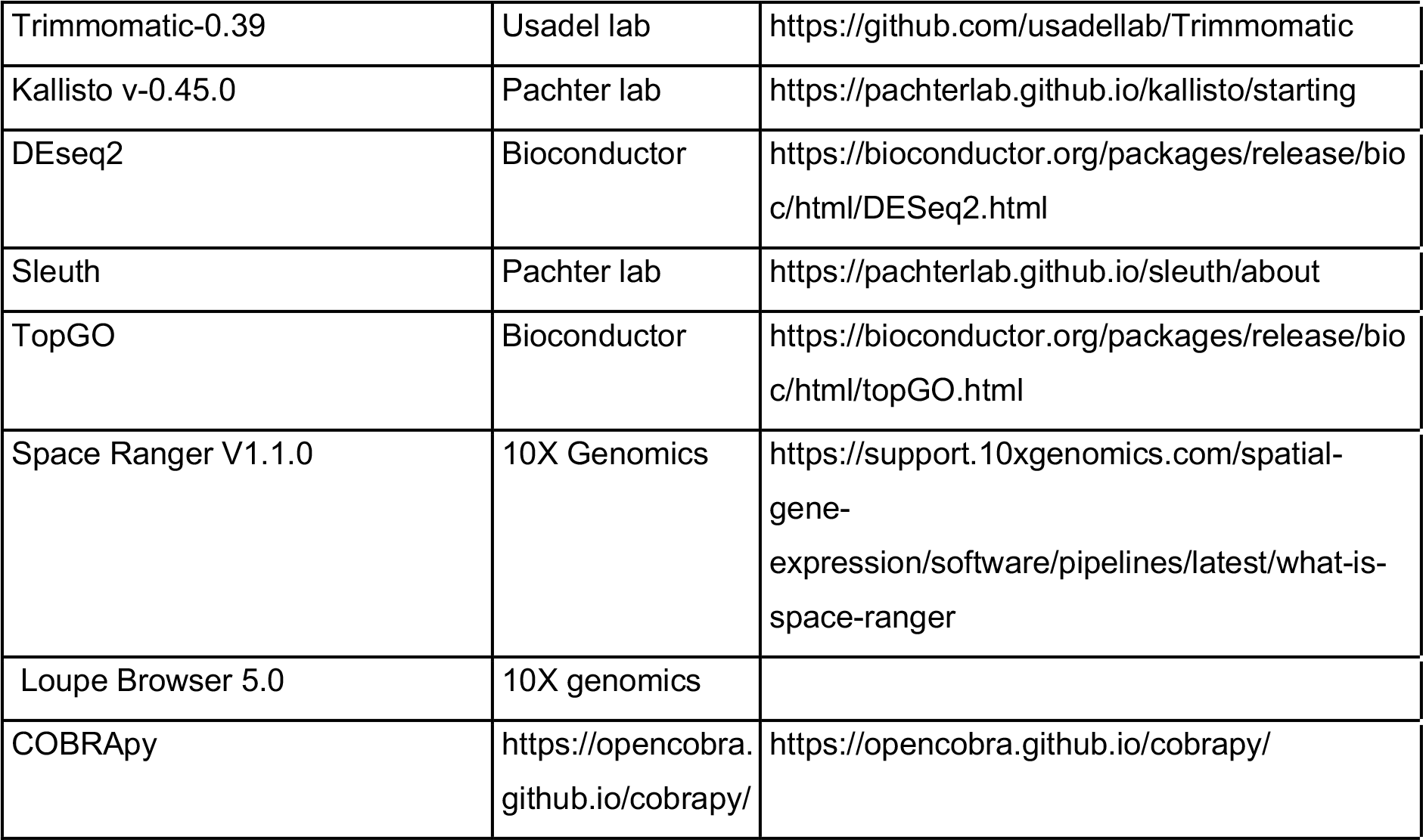
KEY RESOURCES TABLE

## Data and materials availability

Table S10 to S13 (.xlsx) available on Dryad DOI https://doi.org/10.5061/dryad.931zcrjm6 Visium Loupe Browser (.loupe) files Data S1 to S6 available on Dryad DOI https://doi.org/10.5061/dryad.931zcrjm6

All short reads available in the GeneBank Sequence Read Archive (SRA) BioProject: PRJNA774250

## Supplementary Materials

Methods I: Spatial Gene Expression Results I: Spatial Gene Expression Figs. S1 to S7

Tables S1 to S5

Methods II: Flux balance model Results II: Flux balance model Figs. S8 to S10

Tables S6 to S9

Captions for Table S10 to S13 Captions for Data S1 to S6

