## Supplement for "Spatial resolution of an integrated C_4_+CAM photosynthetic metabolism"

### Supplementary Materials

**This PDF file includes:**

- Figs. S1 to S7
- Tables S1 to S7
- Captions for Online Tables S8 to S11
- Captions for Online Data S1 to S6

**Other Supplementary Materials for this manuscript include the following**

- Table S8 to S11 (.xlsx) available on Dryad DOI https://doi.org/10.5061/dryad.931zcrjm6
- Visium loupe files Data S1 to S6 (.loupe) available on Dryad DOI https://doi.org/10.5061/dryad.931zcrjm6
- All short sequencing reads are available in the GeneBank Sequence Read Archive (SRA) BioProject: PRJNA774250
- The scripts used for the data analyses are available on https://github.com/josemovi/Portulaca-spatial-C4-CAM

### Supplementary figures

**Fig. S1. Gene Ontology enrichment across bundle sheath and mesophyll and across watering regimens (extension of fig. 2).**

50 most significant Gene Ontology terms enriched across differentially expressed genes between mesophyll and bundle sheath. Barplots indicate the percentage of genes up-regulated of each GO term across (A) mesophyll and bundle sheath, (B) well-watered and drought, (C) 23h and 7h.

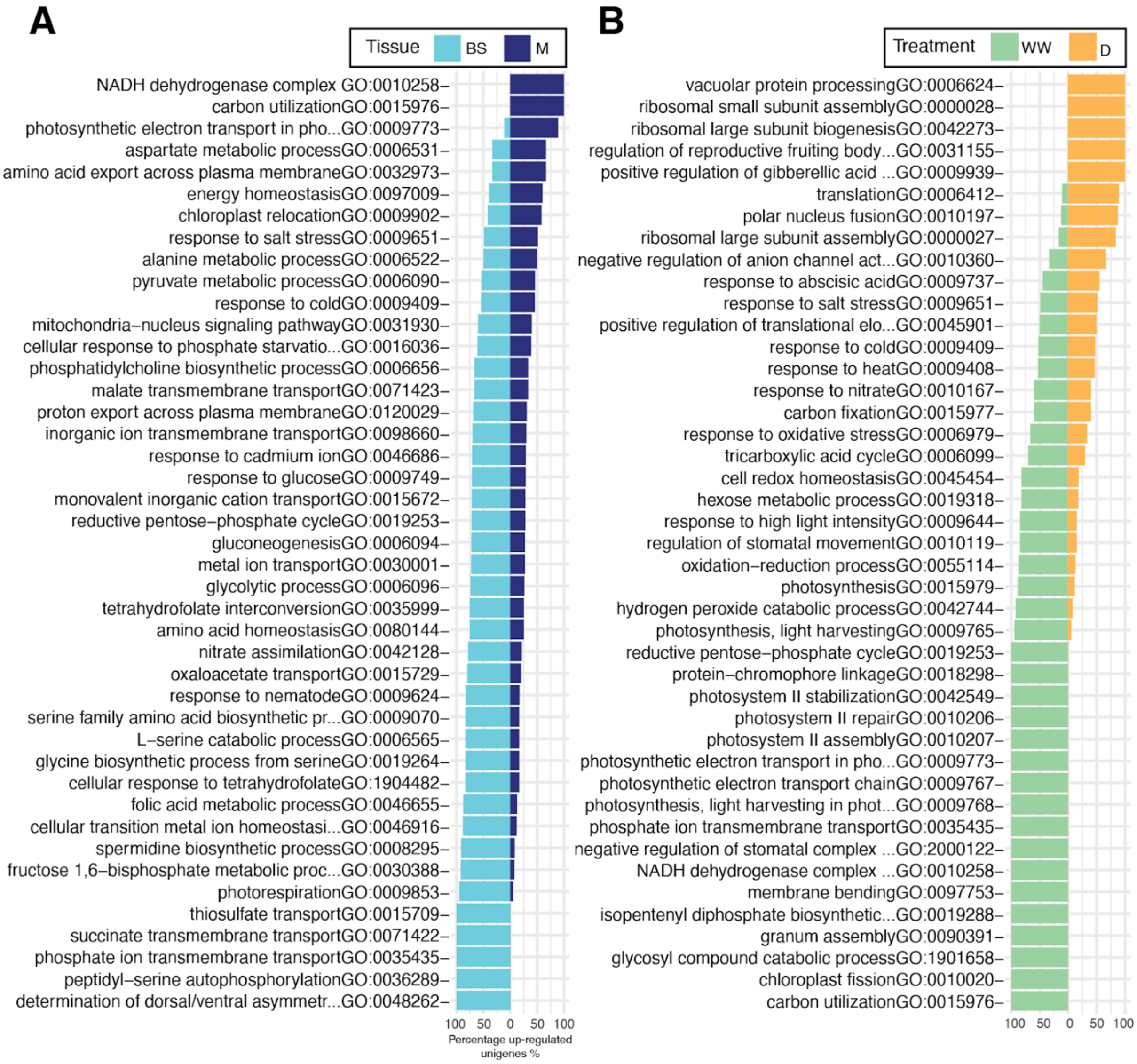

**Figure S2. Figure S2. Differential transcript abundance across cell types and experimental conditions (extended fig. 3)** Differential transcript abundance (measured in log2 fold change, log2FC) of selected genes in mesophyll (M) relative to bundle sheath (BS) tissue (left panel), 07h relative to 23h (middle panel) and drought relative to well-watered (right panel) across LMD samples. Gene colour backgrounds correspond with pathways in the boxes on the top. In all panels, asterisks indicate significant differential expression (*P*adj < 0.05).

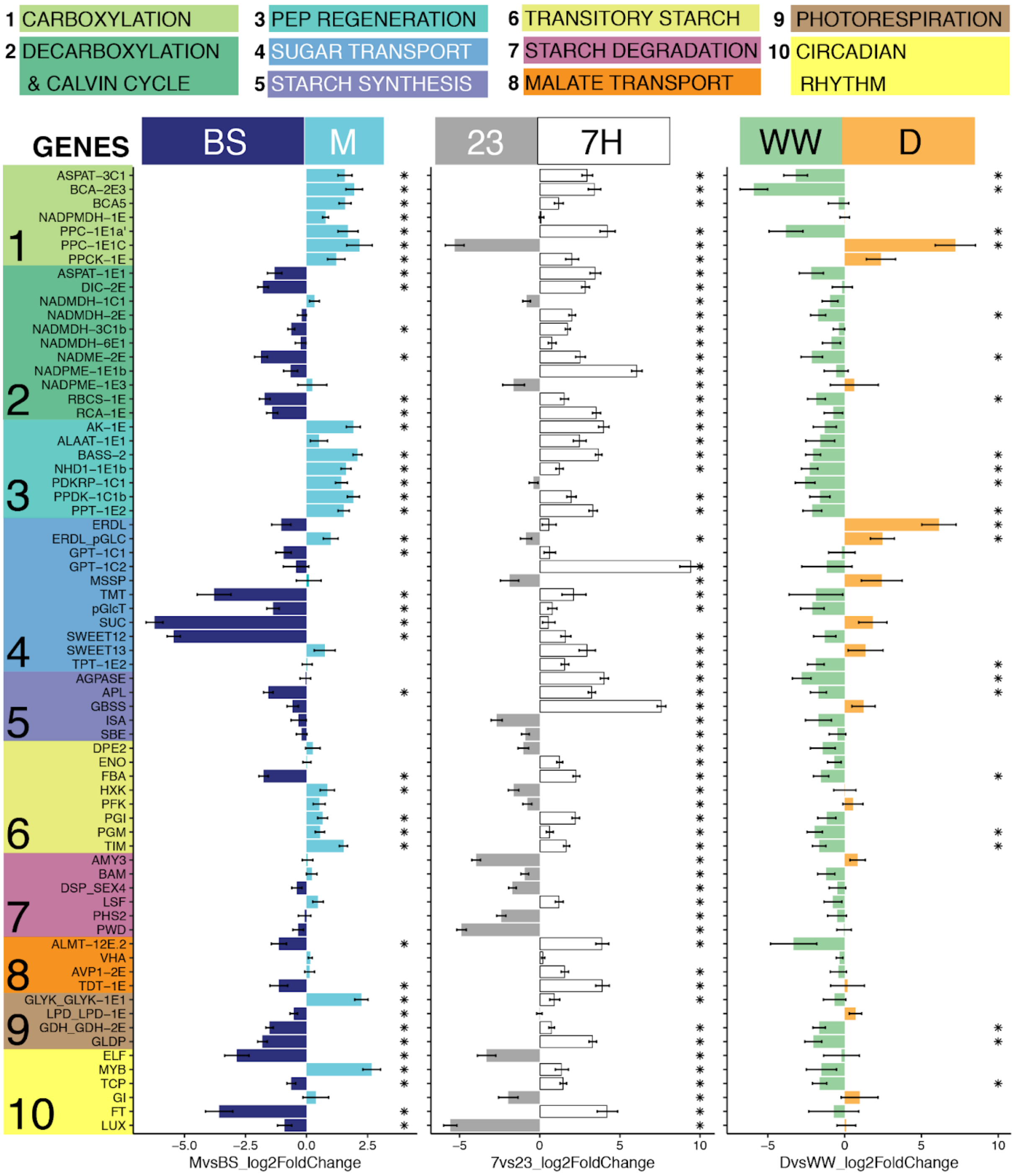

**Figure S3. Figure S3. Effect of watering regimen in each cell type across time points (extension fig. 3)**. Differential transcript abundance in droughted:D relative to well-watered plants:WW (measured in log2 fold change, log2FC). Gene colour backgrounds correspond to pathways in the boxes on the top. Triangles on the left panel represent relative abundance of D mesophyll samples vs WW mesophyll, in 7h samples (white triangles) and in 23h samples (black triangles). Right panel shows the same abundance comparisons using only bundle sheath samples. A triangle in the WW region (negative log2FC, left to the red lines) indicates higher expression during WW, while triangles within the D region (right to the red line) indicate higher expression in D. Asterisks indicate significant differential expression (*P*adj < 0.05)

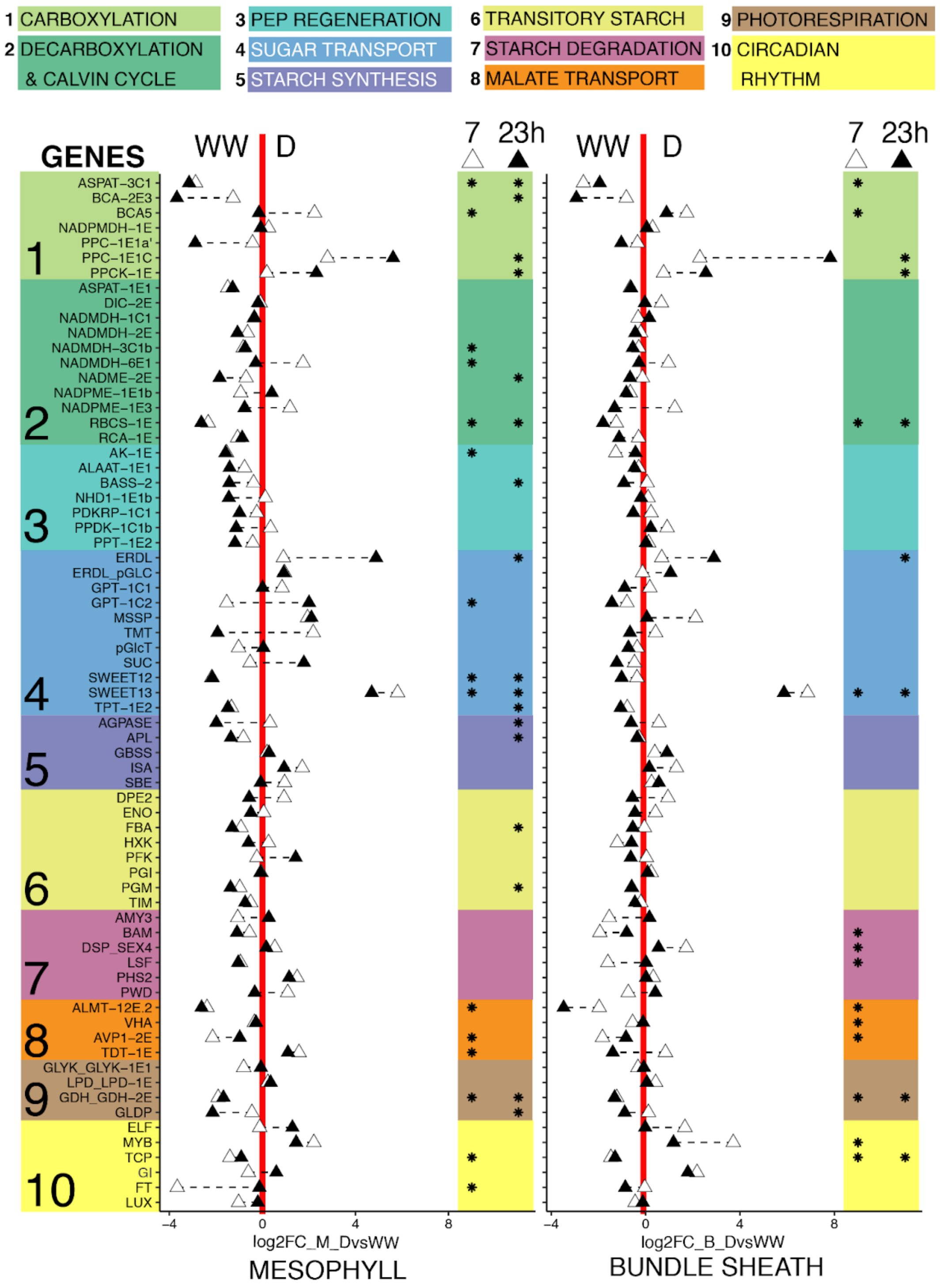

**Fig. S4. Transcription abundance of CCM-related genes and genes with no known role in C4 or CAM (extension of Fig. 4).** A) Transcription abundance of CCM-related genes listed in table S3. B) Genes with no known role in C4 or CAM listed in table S4. Median of transcripts per million (y-axis), across time points (x-axis) in LMD mRNA libraries. Black and red lines indicate mesophyll or bundle sheath, respectively. Plain lines indicate watered and dotted indicate drought. Error bars show the interquartile range of expression.

# A

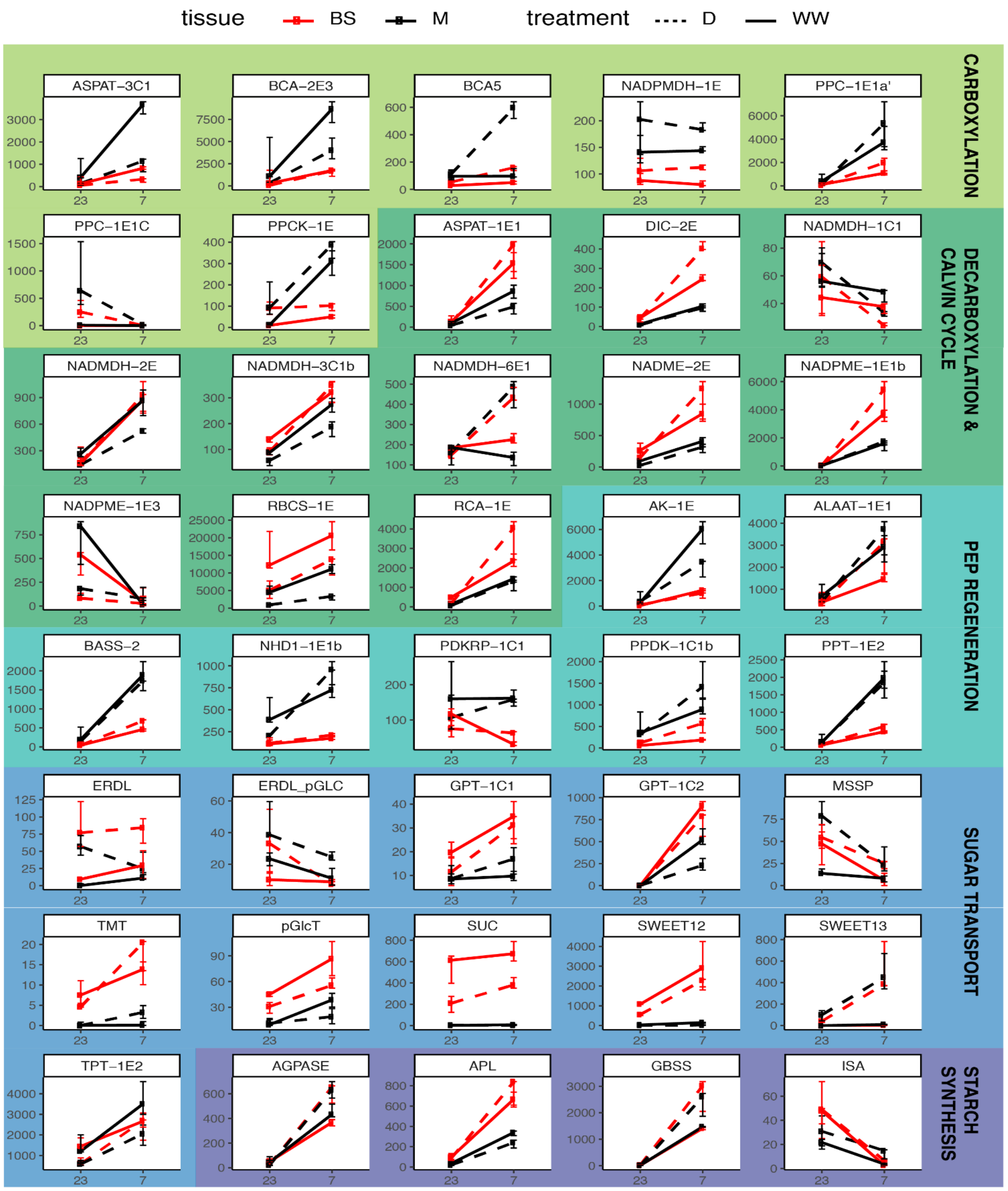

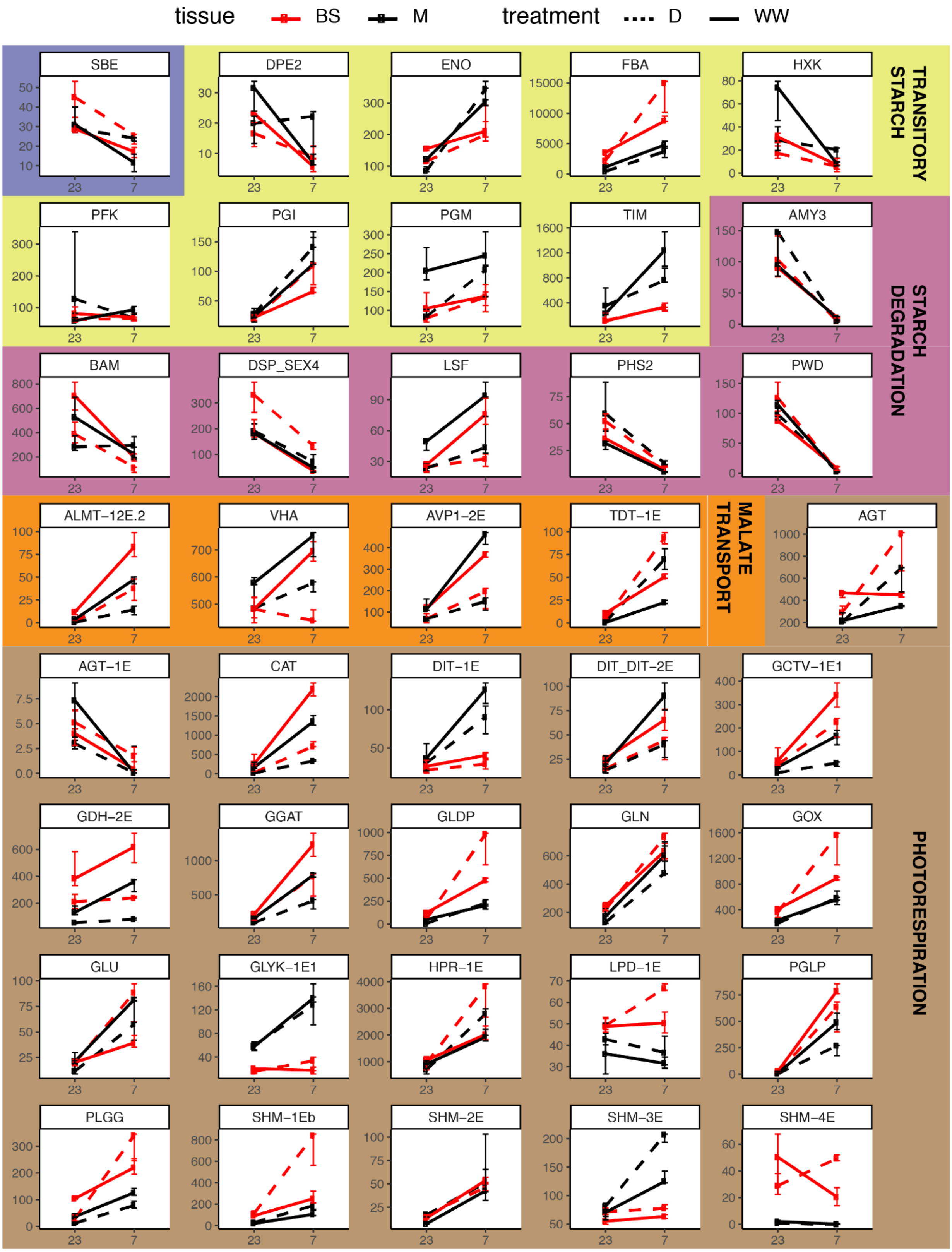
tissue BS M
treatment D WW
SBE
DPE2
ENO
FBA
HXK
TRANSITORY STARCH
PFK
PGI
PGM
TIM
AMY3
STARCH DEGRADATION
BAM
DSP_SEX4
LSF
PHS2
PWD
ALMT−12E.2
VHA
AVP1−2E
TDT−1E
MALATE TRANSPORT
AGT
AGT−1E
CAT
DIT−1E
DIT_DIT−2E
GCTV−1E1
GDH−2E
GGAT
GLDP
GLN
GOX
GLU
GLYK−1E1
HPR−1E
LPD−1E
PGLP
PLGG
SHM−1Eb
SHM−2E
SHM−3E
SHM−4E
PHOTORESPIRATION
23
7

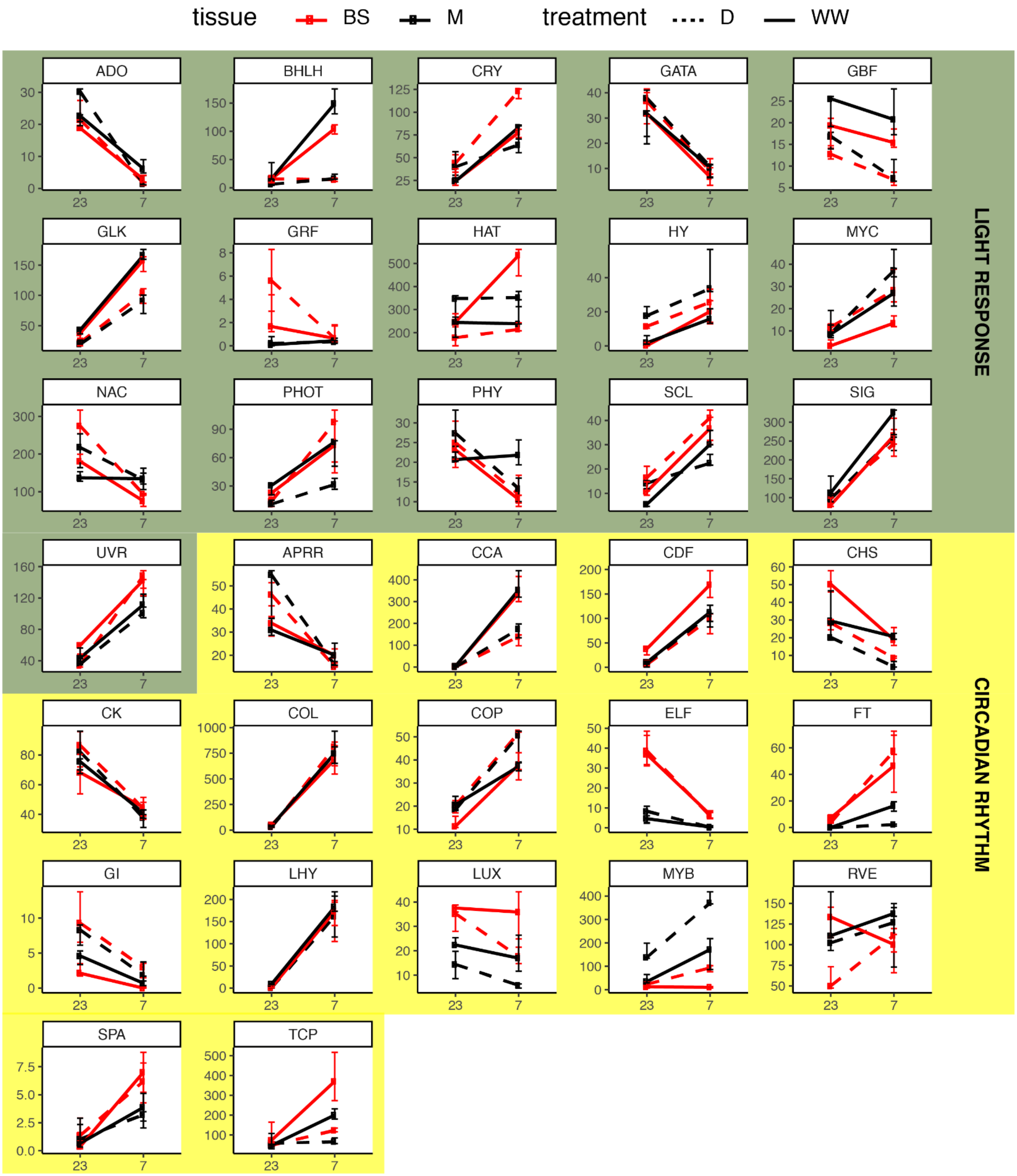
tissue
BS
M
treatment
D
WW
ADO
BHLH
CRY
GATA
GBF
GLK
GRF
HAT
HY
MYC
NAC
PHOT
PHY
SCL
SIG
UVR
APRR
CCA
CDF
CHS
CK
COL
COP
ELF
FT
GI
LHY
LUX
MYB
RVE
SPA
TCP
LIGHT RESPONSE
CIRCADIAN RHYTHM

**B**

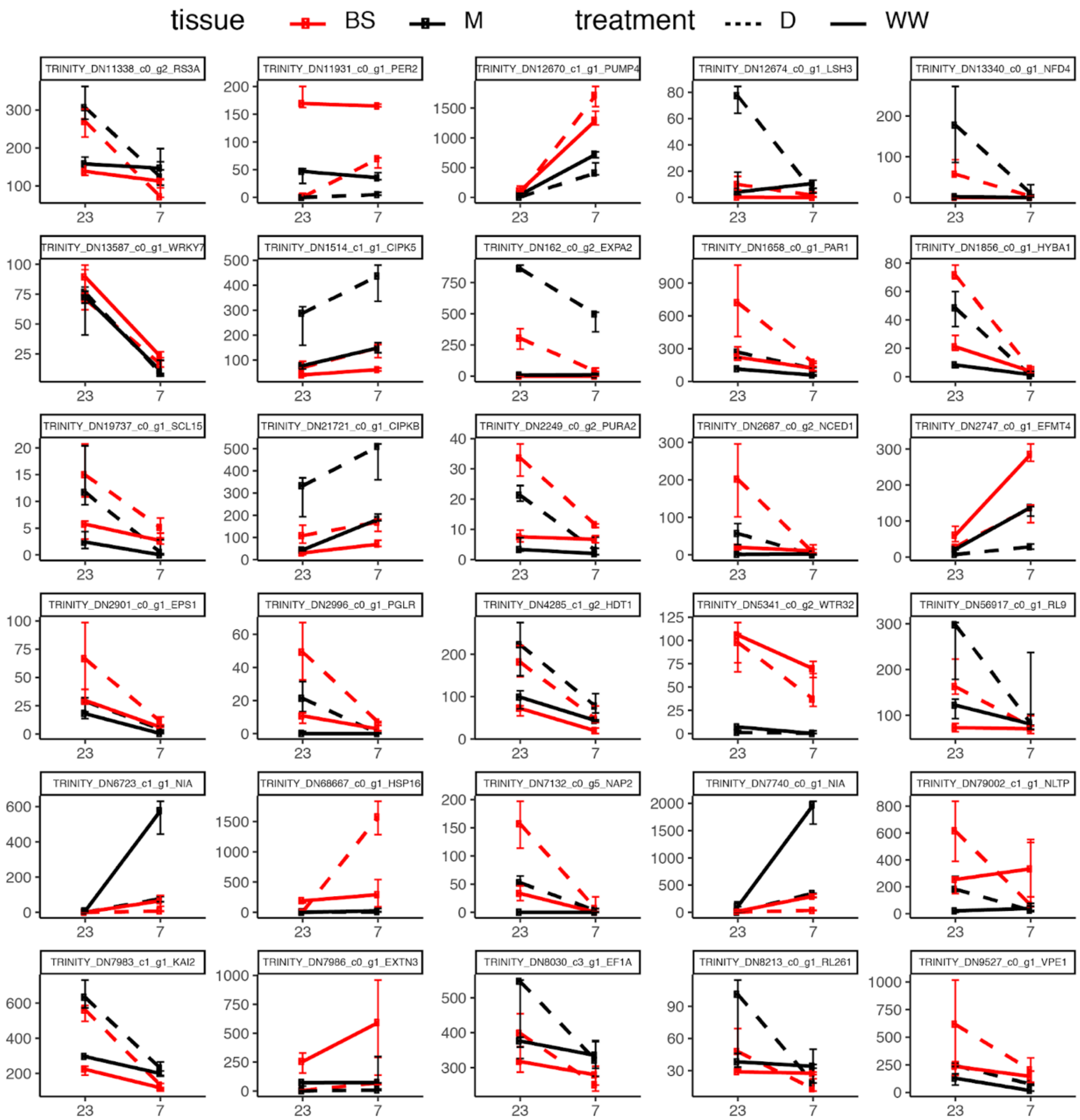

tissue
BS
M
treatment
D
WW
TRINITY_DN11338_c0_g2_RS3A
TRINITY_DN11931_c0_g1_PER2
TRINITY_DN12670_c1_g1_PUMP4
TRINITY_DN12674_c0_g1_LSH3
TRINITY_DN13340_c0_g1_NFD4
TRINITY_DN13587_c0_g1_WRKY7
TRINITY_DN1514_c1_g1_CIPK5
TRINITY_DN162_c0_g2_EXPA2
TRINITY_DN1658_c0_g1_PAR1
TRINITY_DN1856_c0_g1_HYBA1
TRINITY_DN19737_c0_g1_SCL15
TRINITY_DN21721_c0_g1_CIPKB
TRINITY_DN2249_c0_g2_PURA2
TRINITY_DN2687_c0_g2_NCED1
TRINITY_DN2747_c0_g1_EFMT4
TRINITY_DN2901_c0_g1_EPS1
TRINITY_DN2996_c0_g1_PGLR
TRINITY_DN4285_c1_g2_HDT1
TRINITY_DN5341_c0_g2_WTR32
TRINITY_DN56917_c0_g1_RL9
TRINITY_DN6723_c1_g1_NIA
TRINITY_DN68667_c0_g1_HSP16
TRINITY_DN7132_c0_g5_NAP2
TRINITY_DN7740_c0_g1_NIA
TRINITY_DN79002_c1_g1_NLTP
TRINITY_DN7983_c1_g1_KAI2
TRINITY_DN7986_c0_g1_EXTN3
TRINITY_DN8030_c3_g1_EF1A
TRINITY_DN8213_c0_g1_RL261
TRINITY_DN9527_c0_g1_VPE1

**Figure S5. Visium spatial gene expression (extended fig. 5)**. The first row shows frozen leaf specimens at the moment of cryo-sectioning. The second row shows microphotographs of leaf paradermal sections under bright field (left) and K-means clustering of total gene expression (right). Successive rows show abundance of the main CCM-related genes using the 10x Genomics Visium platform. K-means clustering of sampling spots corresponds to bundle sheath (BS, dark blue), mesophyll (M, light blue), and water storage (WS, orange) tissues; abundances are shown relative to their observed unique molecule index (UMI) ranges.

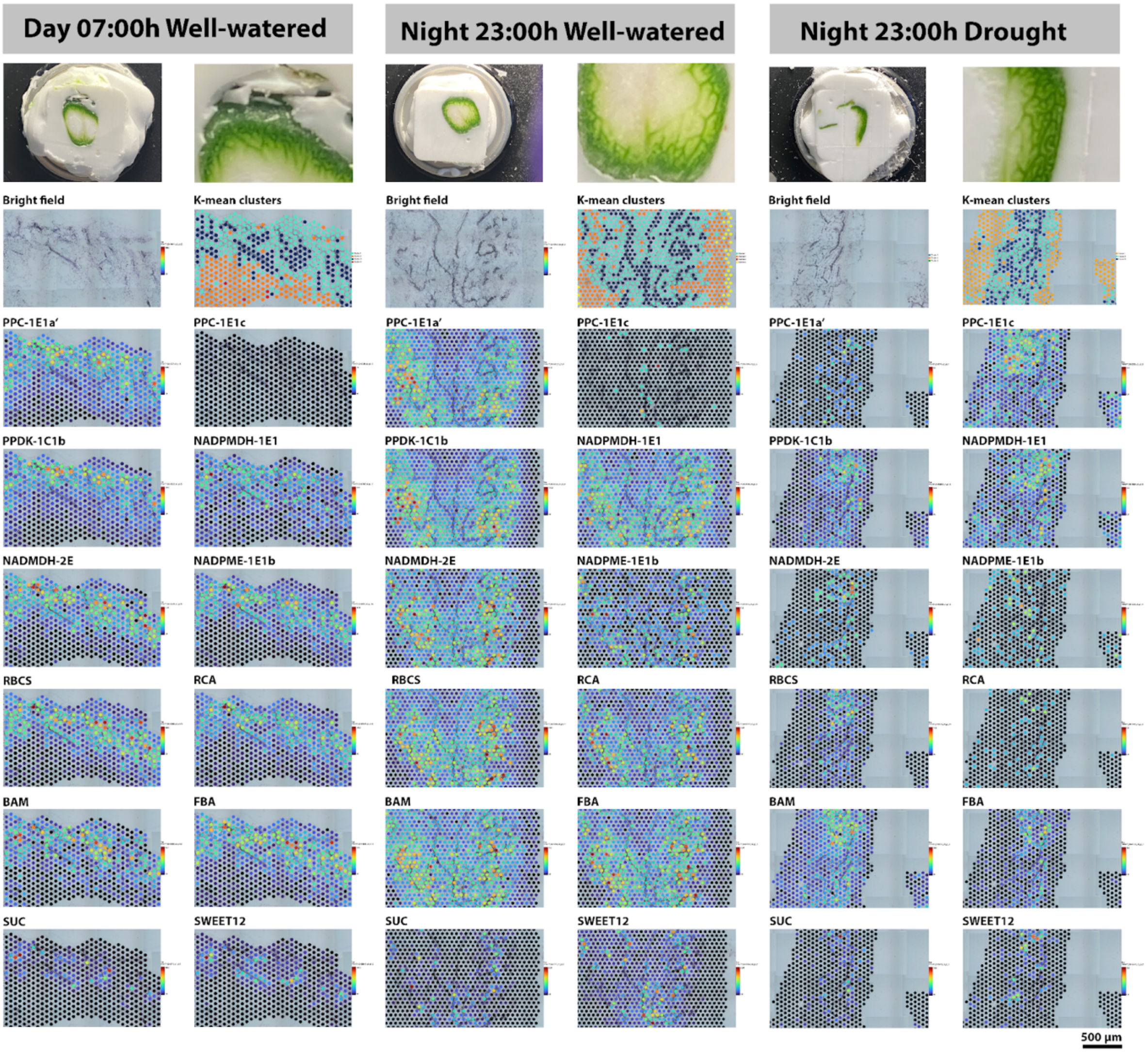

**Fig. S6. Correlations between predicted enzymatic fluxes and estimated gene expression in mesophyll and bundle sheath samples**. Pearson correlation between z-score normalized (mean set to 0, SD set to 1) pFBA results and transcript abundance. The transcript abundances from different orthologs used in the same biochemical reactions were added together to be compared with the pFBA results. 23_BS_D: nighttime flux in bundle sheath under drought; 23_BS_W: nighttime flux in bundle sheath under well-water; 23_M_D: nighttime flux in mesophyll under drought; 23_M_W: nighttime flux in mesophyll under well-water; 7_BS_D: daytime flux in bundle sheath under drought; 7_BS_W: daytime flux in bundle sheath under well-water; 7_M_D: daytime flux in mesophyll under drought; 7_M_W: daytime flux in mesophyll under well-water;

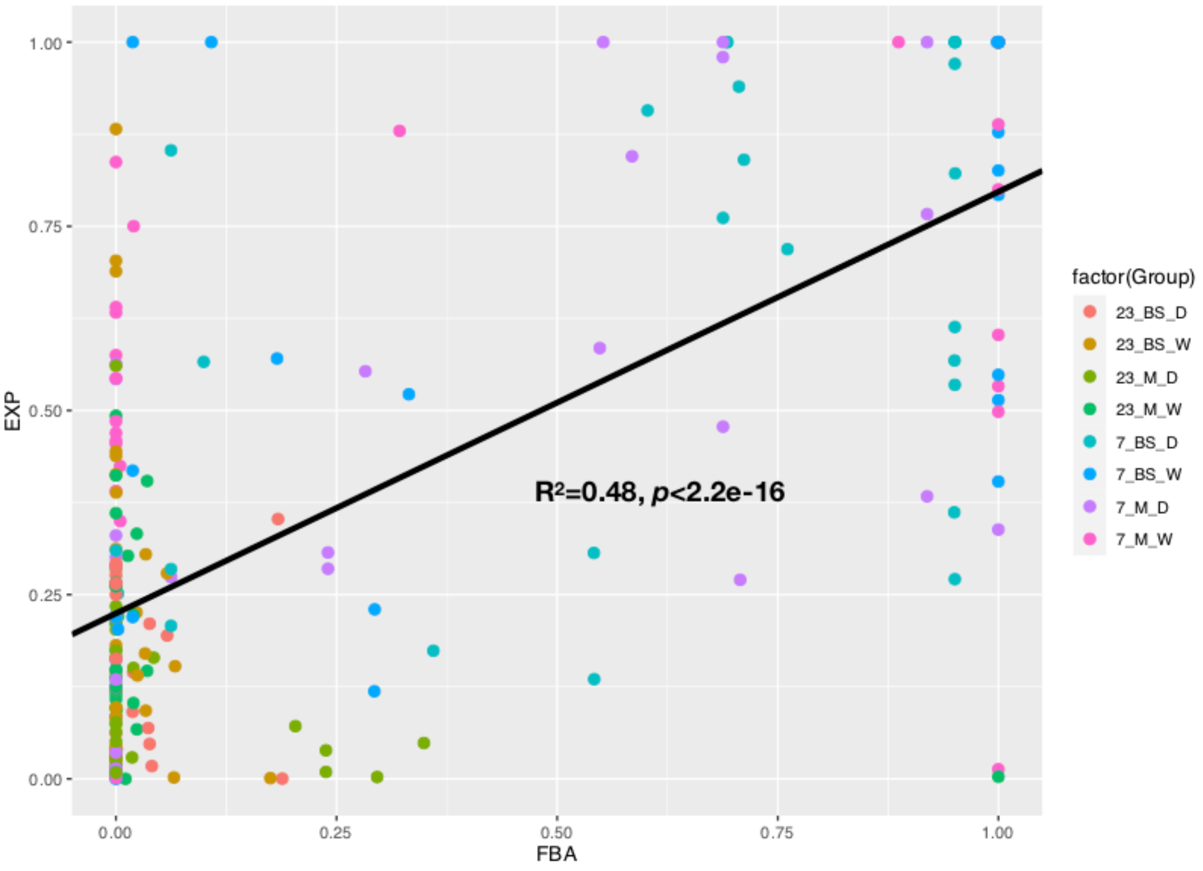

**Fig. S7. Flux variability analysis (FVA) of mesophyll and bundle sheath with diel variation and under drought and well-water conditions (related to Fig. 7).** Flux variability analysis (FVA) of mesophyll and bundle sheath with diel variation and under drought and well-water conditions. Daytime: orange arrows and white background; nighttime: blue arrows and grey background. Reactions that occur during the day-night transition are shown with striped arrows. M: mesophyll; BS: bundle sheath.Black numbers: well-water condition (300 ppm CO2 concentration at daytime and and no limit for nighttime); red number: drought condition (70 ppm CO2 concentration at daytime and no limit for nighttime).

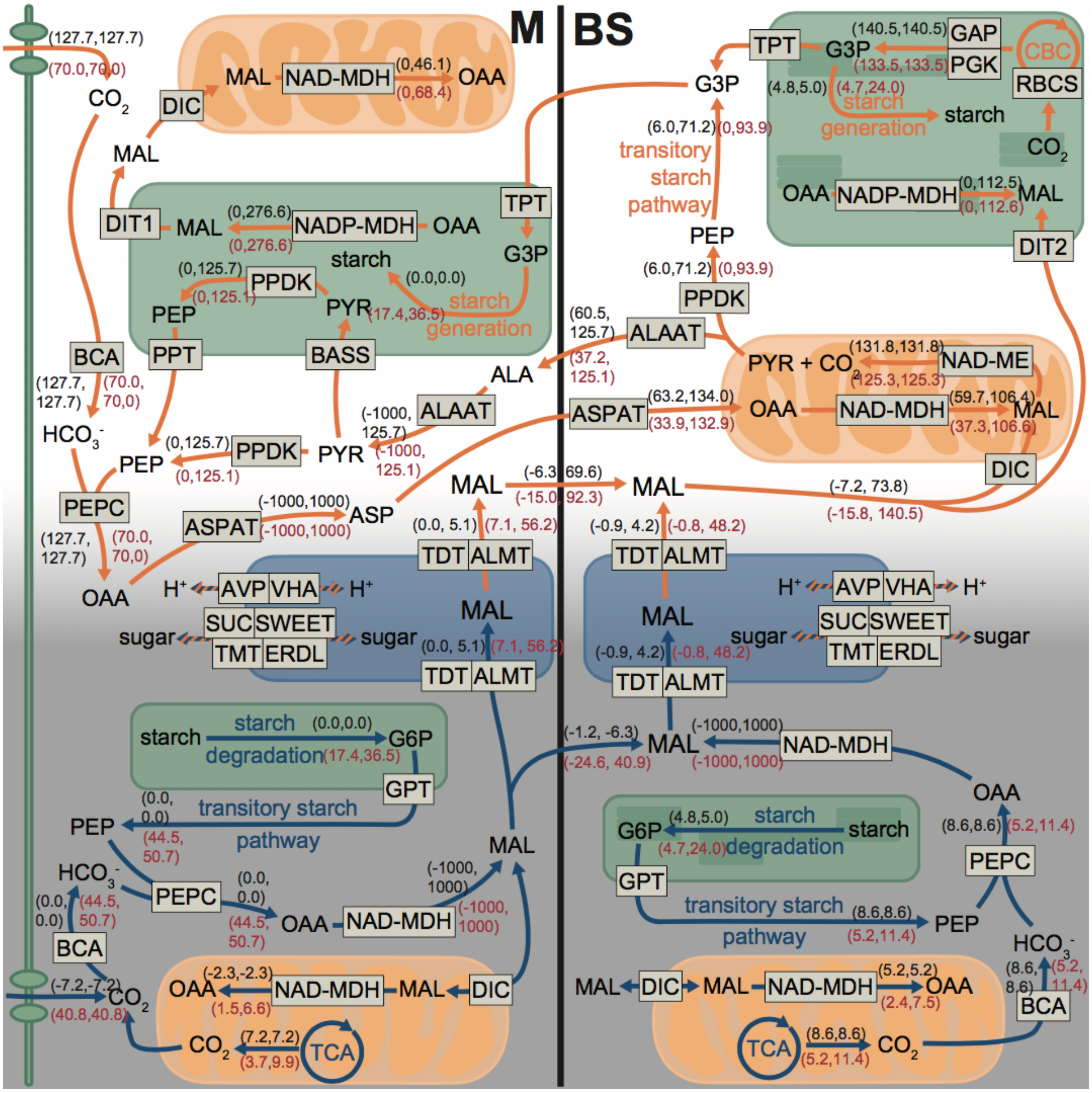

### Supplementary tables

**Table S1. (Related to Fig. 1A).**

Leaf titratable acidity analysis results. Microequivalents H+ (μEq H+) per gram fresh mass was calculated as volume titrant (μL) × titrant molarity (M) / tissue mass (g).

| Sample_id | Treatment | Time (24h) | μEq H+ |
|---|---|---|---|
| POL1-D-7 | drought | 7h:00 | 76.22 |
| POL1-D-7 | drought | 7h:00 | 89.52 |
| POL2-D-7 | drought | 7h:00 | 84.13 |
| POL2-D-7 | drought | 7h:00 | 71.11 |
| POL1-D-19 | drought | 19:00:00 | 27.12 |
| POL1-D-19 | drought | 19:00:00 | 22.26 |
| POL2-D-19 | drought | 19:00:00 | 41.24 |
| POL2-D-19 | drought | 19:00:00 | 11.05 |
| POL1-W-7 | watered | 7h:00 | 24.32 |
| POL1-W-7 | watered | 7h:00 | 15.63 |
| POL2-W-7 | watered | 7h:00 | 22.32 |
| POL2-W-7 | watered | 7h:00 | 16.73 |
| POL1-W-19 | watered | 19:00:00 | 23.41 |
| POL1-W-19 | watered | 19:00:00 | 25.96 |
| POL2-W-19 | watered | 19:00:00 | 16.75 |
| POL2-W-19 | watered | 19:00:00 | 16.97 |

**Table S2 (related to Fig. 2).**

LMD-RNA sequencing and read mapping statistics. BS: bundle-sheath cells, M: mesophyll cells, n_: number of reads.T: time; Ind: Plant individual; n_raw: number of raw reads; n_filtered: number of reads after filtering; n_aligned: number of aligned read to the transcriptome; %_al: percentage of reads aligned to the transcriptome.

| Library_id | T | Tissue | Treatment | Ind | n_raw | n_filtered | n_aligned | %_al |
|---|---|---|---|---|---|---|---|---|
| 22-A_PO1-D-7-M | 7h | BS | Drought | PO1 | 37,306,803 | 32,891,937 | 25,374,659 | 77.10 |
| 5-2-A_PO2-D-7-BS | 7h | BS | Drought | PO2 | 36,651,612 | 32,289,322 | 23,469,722 | 72.70 |
| 5-A_PO2-D-7-BS | 7h | BS | Drought | PO2 | 32,587,781 | 28,351,390 | 20,341,689 | 71.70 |
| 10-B_PO2-WW-23-BS | 7h | BS | Watered | PO1 | 39,734,057 | 23,311,582 | 17,349,512 | 74.40 |
| 12-B_PO1-WW-7-BS | 7h | BS | Watered | PO1 | 34,351,505 | 23,369,981 | 17,052,138 | 73.00 |
| 4-12-B_PO1-WW-7-BS | 7h | BS | Watered | PO1 | 32,181,954 | 20,082,323 | 14,527,796 | 72.30 |
| 19-B_PO2-WW-7-BS | 7h | BS | Watered | PO2 | 37,865,848 | 19,888,745 | 14,497,329 | 72.90 |
| 24-A_PO1-D-23-M | 7h | M | Drought | PO1 | 35,708,771 | 27,081,395 | 22,110,992 | 81.60 |
| 4-2-A_PO2-D-7-M | 7h | M | Drought | PO2 | 31,310,261 | 27,982,988 | 21,524,928 | 76.90 |
| 6-2-A_PO2-D-7-M | 7h | M | Drought | PO2 | 35,647,657 | 31,989,325 | 24,098,407 | 75.30 |
| 13-B_PO1-WW-7-M | 7h | M | Watered | PO1 | 34,519,270 | 22,373,676 | 16,661,642 | 74.50 |
| 15-B_PO1-WW-7-M | 7h | M | Watered | PO1 | 41,545,559 | 29,425,690 | 21,639,546 | 73.50 |
| 21-A_PO1-D-7-BS | 7h | M | Watered | PO2 | 39,268,580 | 34,697,032 | 22,238,196 | 64.10 |
| 26-A_PO1-D-23-M | 23h | BS | Drought | PO1 | 38,373,081 | 32,491,941 | 26,358,868 | 81.10 |
| 7-C_PO1-D-23-BS | 23h | BS | Drought | PO1 | 35,134,577 | 15,276,839 | 10,700,178 | 70.00 |
| 15-A_PO2-D-23-BS | 23h | BS | Drought | PO2 | 38,199,397 | 29,299,221 | 21,694,673 | 74.00 |
| 7-A_PO2-D-23-BS | 23h | BS | Drought | PO2 | 35,729,969 | 30,921,888 | 23,884,264 | 77.20 |
| 3-C_PO1-WW-23-BS | 23h | BS | Watered | PO1 | 39,224,405 | 17,905,370 | 13,410,110 | 74.90 |
| 1-12-B_PO1-WW-7-BS | 23h | BS | Watered | PO2 | 33,572,474 | 19,664,420 | 14,754,741 | 75.00 |
| 8-B_PO2-WW-23-BS | 23h | BS | Watered | PO2 | 38,190,910 | 20,875,273 | 15,495,580 | 74.20 |
| 25-A_PO1-D-23-BS | 23h | M | Drought | PO1 | 35,543,707 | 31,380,508 | 24,151,852 | 77.00 |
| 2-C_PO1-WW-23-M | 23h | M | Drought | PO1 | 35,114,605 | 12,695,318 | 10,070,151 | 79.30 |
| 8-C_PO1-D-23-M | 23h | M | Drought | PO1 | 48,944,420 | 19,757,250 | 15,312,762 | 77.50 |
| 16-A_PO2-D-23-M | 23h | M | Drought | PO2 | 40,444,993 | 34,196,837 | 27,231,638 | 79.60 |
| 8-A_PO2-D-23-M | 23h | M | Drought | PO2 | 35,841,757 | 30,860,105 | 24,858,317 | 80.60 |
| 20-B_PO2-WW-7-M | 23h | M | Watered | PO1 | 39,592,141 | 22,769,276 | 16,909,912 | 74.30 |
| 11-B_PO2-WW-23-M | 23h | M | Watered | PO2 | 44,046,759 | 28,007,152 | 21,373,120 | 76.30 |
| 9-B_PO2-WW-23-M | 23h | M | Watered | PO2 | 43,071,551 | 24,150,596 | 18,659,731 | 77.30 |

**Table S3. (related to Figs. 3-4-5-S2-S3)** Annotation of selected genes with a role or potential role in CCM.

| Phylo-annotation | Uniprot_gene | Description | unigenes | Pathway |
|---|---|---|---|---|
| ASPAT-3C1 | AATC_DAUCA | Aspartate aminotransferase, cytoplasmic | TRINITY_DN346_c1_g3 | Carboxylation |
| BCA-2E3 | | Beta carbonic anhydrase | TRINITY_DN889_c1_g2 | Carboxylation |
| BCA5 | BCA5_ARATH | Beta carbonic anhydrase 5, chloroplastic | TRINITY_DN8226_c0_g1 | Carboxylation |
| NADPMDH-1E | MDHP_MESCR | Malate dehydrogenase [NADP], chloroplastic | TRINITY_DN11597_c0_g1 | Carboxylation |
| PPC-1E1a' | CAPP_AMAHP | Phosphoenolpyruvate carboxylase | TRINITY_DN1747_c2_g1 | Carboxylation |
| PPC-1E1C | CAPP2_ARATH | Phosphoenolpyruvate carboxylase 2 | TRINITY_DN3235_c0_g3 | Carboxylation |
| PPCK-1E | PPCK1_ARATH | Phosphoenolpyruvate carboxylase kinase 1 | TRINITY_DN4567_c3_g1 | Decarboxylation |
| ASPAT-1E1 | | Alanine aminotransferase 2, mitochondrial | TRINITY_DN18480_c0_g1 | Decarboxylation |
| DIC-2E | PUMP5_ARATH | Mitochondrial uncoupling protein 5 | TRINITY_DN1178_c2_g2 | Decarboxylation |
| NADMDH-1C1 | MDHP_ARATH | Malate dehydrogenase, chloroplastic | TRINITY_DN2897_c0_g1 | Decarboxylation |
| NADMDH-2E | | Malate dehydrogenase | TRINITY_DN12466_c1_g1 | Decarboxylation |
| NADMDH-3C1b | MDHG_CUCSA | Malate dehydrogenase, glyoxysomal | TRINITY_DN379_c2_g1 | Decarboxylation |
| NADMDH-6E1 | MDHC_BETVU | Malate dehydrogenase, cytoplasmic | TRINITY_DN70366_c1_g1 | Decarboxylation |
| NADME-2E | MAON_SOLTU | NAD-dependent malic enzyme 59 kDa isoform, mitochondrial | TRINITY_DN17872_c1_g1 | Decarboxylation |
| NADPME-1E1b | MAOC_FLAPR | NADP-dependent malic enzyme, chloroplastic | TRINITY_DN1253_c0_g1 | Decarboxylation |
| NADPME-1E3 | MAOX_VITVI | NADP-dependent malic enzyme | TRINITY_DN9969_c0_g1 | Decarboxylation |
| RBCS-1E | | Ribulose bisphosphate carboxylase small chain, chloroplastic | TRINITY_DN6389_c0_g1 | Calvin cycle |
| RCA-1E | | Ribulose bisphosphate carboxylase/oxygenase activase, chloroplastic | TRINITY_DN17967_c2_g1 | Calvin cycle |
| AK-1E | KAD2_ARATH | Adenylate kinase 2, chloroplastic | TRINITY_DN1795_c0_g4 | PEP generation |
| ALAAT-1E1 | ALAT2_ARATH | Alanine aminotransferase 2, mitochondrial | TRINITY_DN8707_c0_g1 | PEP generation |
| BASS-2 | BASS2_ARATH | Sodium/pyruvate cotransporter BASS2, chloroplastic | TRINITY_DN6617_c4_g1 | PEP generation |
| NHD1-1E1b | NHD1_ARATH | Sodium/proton antiporter 1 | TRINITY_DN1372_c2_g1 | PEP generation |
| PDKRP-1C1 | PDRP1_ORYSI | Probable pyruvate, phosphate dikinase regulatory protein, chloroplastic | TRINITY_DN7731_c0_g1 | PEP generation |
| PPDK-1C1b | PPDK_MESCR | Pyruvate, phosphate dikinase, chloroplastic | TRINITY_DN4872_c3_g1 | PEP generation |
| PPT-1E2 | PPT2_ORYSJ | Phosphoenolpyruvate/phosphate translocator 2, chloroplastic | TRINITY_DN13020_c1_g1 | PEP generation |
| ERDL | EDL16_ARATH | Sugar transporter ERD6-like 16 | TRINITY_DN1122_c0_g1 | Starch/sugar transport |
| ERDL_pGLC | ERDL7_ARATH | Sugar transporter ERD6-like 7 | TRINITY_DN2349_c0_g1 | Starch/sugar transport |
| GPT-1C1 | GPT2_ARATH | Glucose-6-phosphate/phosphate translocator 2, chloroplastic | TRINITY_DN6031_c0_g1 | Starch/sugar transport |
| GPT-1C2 | GPT2_ARATH | Glucose-6-phosphate/phosphate translocator 2, chloroplastic | TRINITY_DN6322_c1_g1 | Starch/sugar transport |
| MSSP | MSSP2_ARATH | Monosaccharide-sensing protein 2 | TRINITY_DN17666_c0_g1 | Starch/sugar transport |
| TMT | MSSP2_ARATH | Monosaccharide-sensing protein 2 | TRINITY_DN6869_c0_g2 | Starch/sugar transport |
| pGlcT | PLST4_ARATH | Plastidic glucose transporter 4 | TRINITY_DN7812_c0_g2 | Starch/sugar transport |
| SUC | SUT_SPIOL | Sucrose transport protein | TRINITY_DN6710_c1_g1 | Starch/sugar transport |
| SWEET12 | SWT12_ARATH | Bidirectional sugar transporter SWEET12 | TRINITY_DN19841_c0_g1 | Starch/sugar transport |
| SWEET13 | SWT13_ARATH | Bidirectional sugar transporter SWEET13 | TRINITY_DN8938_c4_g2 | Starch/sugar transport |
| TPT-1E2 | TPT_SPIOL | Triose phosphate/phosphate translocator, chloroplastic | TRINITY_DN1249_c1_g1 | Starch/sugar transport |
| AGPASE | GLGS2_VICFA | Glucose-1-phosphate adenylyltransferase small subunit 2, chloroplastic | TRINITY_DN7789_c1_g1 | Starch synthesis |
| APL | GLGL1_ARATH | Glucose-1-phosphate adenylyltransferase large subunit 1, chloroplastic | TRINITY_DN4352_c0_g1 | Starch synthesis |
| GBSS | SSG1_MANES | Granule-bound starch synthase 1, chloroplastic/amyloplastic | TRINITY_DN11204_c1_g1 | Starch synthesis |
| ISA | ISOA1_ARATH | Isoamylase 1, chloroplastic | TRINITY_DN4229_c0_g1 | Starch synthesis |
| SBE | GLGB1_PEA | 1,4-alpha-glucan-branching enzyme 1, chloroplastic/amyloplastic | TRINITY_DN2618_c0_g1 | Starch synthesis |
| DPE2 | DPE2_ARATH | 4-alpha-glucanotransferase DPE2 | TRINITY_DN6357_c0_g1 | Transitory starch pathway |

| | | | | |
|---|---|---|---|---|
| ENO | ENO_MESCR | Enolase | TRINITY_DN6068_c1_g2 | Transitory starch pathway |
| FBA | ALFP_ORYSJ | Fructose-bisphosphate aldolase, chloroplastic | TRINITY_DN1271_c0_g1 | Transitory starch pathway |
| HXK | HXK2_ORYSJ | Hexokinase-2 | TRINITY_DN7391_c0_g1 | Transitory starch pathway |
| PFK | PFKA6_ARATH | ATP-dependent 6-phosphofructokinase 6 | TRINITY_DN8265_c0_g1 | Transitory starch pathway |
| PGI | | Glucose-6-phosphate isomerase | TRINITY_DN1518_c1_g1 | Transitory starch pathway |
| PGM | PGMC_MESCR | Phosphoglucomutase, cytoplasmic | TRINITY_DN23625_c0_g1 | Transitory starch pathway |
| TIM | TPIC_SPIOL | Triosephosphate isomerase, chloroplastic | TRINITY_DN4168_c0_g2 | Transitory starch pathway |
| AMY3 | AMY3_ARATH | Alpha-amylase 3, chloroplastic | TRINITY_DN2216_c0_g1 | Starch degradation |
| BAM | BAM3_ARATH | Beta-amylase 3, chloroplastic | TRINITY_DN1866_c0_g1 | Starch degradation |
| DSP_SEX4 | DSP4_CASSA | Phosphoglucan phosphatase DSP4, amyloplastic | TRINITY_DN12877_c0_g1 | Starch degradation |
| LSF | LSF2_ARATH | Phosphoglucan phosphatase LSF2, chloroplastic | TRINITY_DN2348_c0_g1 | Starch degradation |
| PHS2 | PHSL2_SOLTU | Alpha-1,4 glucan phosphorylase L-2 isozyme, chloroplastic/amyloplastic | TRINITY_DN12375_c0_g1 | Starch degradation |
| PWD | GWD1_CITRE | Alpha-glucan water dikinase, chloroplastic | TRINITY_DN3424_c0_g1 | Starch degradation |
| ALMT-12E.2 | ALMTC_ARATH | Aluminum-activated malate transporter 12 | TRINITY_DN10391_c0_g2 | Metabolite transport |
| VHA | | V-type proton ATPase subunit | TRINITY_DN16658_c0_g1 | Metabolite transport |
| AVP1-2E | | Pyrophosphate-energized vacuolar membrane proton pump | TRINITY_DN1954_c0_g1 | Metabolite transport |
| TDT-1E | TDT_ARATH | Tonoplast dicarboxylate transporter | TRINITY_DN161_c0_g1 | Metabolite transport |
| AGT | SGAT_ARATH | Serine--glyoxylate aminotransferase | TRINITY_DN6907_c0_g1 | Photorespiration |
| AGT-1E | AGT23_ARATH | Alanine--glyoxylate aminotransferase 2 homolog 3, mitochondrial | TRINITY_DN3579_c0_g1 | Photorespiration |
| CAT | CATA2_GOSHI | Catalase isozyme 2 | TRINITY_DN193_c1_g1 | Photorespiration |
| DIT-1E | DIT1_SPIOL | Dicarboxylate transporter 1, chloroplastic | TRINITY_DN2503_c0_g1 | Photorespiration |
| DIT-2E | DIT2_SPIOL | Dicarboxylate transporter 2, chloroplastic | TRINITY_DN4387_c0_g2 | Photorespiration |
| GCTV-1E1 | GCST_MESCR | Aminomethyltransferase, mitochondrial | TRINITY_DN2504_c1_g1 | Photorespiration |
| GDH-2E | GCSH_MESCR | Glycine cleavage system H protein, mitochondrial | TRINITY_DN6744_c1_g1 | Photorespiration |
| GGAT | | Glutamate--glyoxylate aminotransferase | TRINITY_DN2102_c0_g1 | Photorespiration |
| GLDP | GCSPA_FLAPR | Glycine dehydrogenase (decarboxylating) A, mitochondrial | TRINITY_DN6006_c0_g1 | Photorespiration |
| GLN | GLNA1_LOTJA | Glutamine synthetase cytosolic isozyme | TRINITY_DN4141_c1_g1 | Photorespiration |
| GOX | GOX_SPIOL | Peroxisomal (S)-2-hydroxy-acid oxidase | TRINITY_DN3751_c0_g1 | Photorespiration |
| GLU | GLTB_SPIOL | Ferredoxin-dependent glutamate synthase, chloroplastic | TRINITY_DN425_c0_g1 | Photorespiration |
| GLYK-1E1 | GLYK_ARATH | D-glycerate 3-kinase, chloroplastic | TRINITY_DN2228_c0_g1 | Photorespiration |
| HPR-1E | HPR1_ARATH | Glycerate dehydrogenase HPR, peroxisomal | TRINITY_DN7689_c0_g3 | Photorespiration |
| LPD-1E | DLDH1_ARATH | Dihydrolipoyl dehydrogenase 1, mitochondrial | TRINITY_DN7076_c1_g1 | Photorespiration |
| PGLP | PGP1A_ARATH | Phosphoglycolate phosphatase 1A, chloroplastic | TRINITY_DN17754_c0_g1 | Photorespiration |
| PLGG | | Plastidal glycolate/glycerate translocator | TRINITY_DN10229_c0_g2 | Photorespiration |
| SHM-1Eb | GLYM_SOLTU | Serine hydroxymethyltransferase, mitochondrial | TRINITY_DN302_c1_g2 | Photorespiration |
| SHM-2E | GLYC6_ARATH | Serine hydroxymethyltransferase 6 | TRINITY_DN1750_c0_g1 | Photorespiration |
| SHM-3E | | Serine hydroxymethyltransferase | TRINITY_DN3209_c0_g1 | Photorespiration |
| SHM-4E | GLYP3_ARATH | Serine hydroxymethyltransferase 3, chloroplastic | TRINITY_DN1307_c0_g1 | Photorespiration |
| ADO | ADO1_ARATH | Adagio protein 1 | TRINITY_DN3633_c0_g1 | Light response |
| BHLH | BH062_ARATH | Transcription factor bHLH62 | TRINITY_DN360_c1_g1 | Light response |
| CRY | CRY1_ARATH | Cryptochrome-1 | TRINITY_DN2794_c2_g1 | Light response |
| GATA | | Putative GATA transcription factor | TRINITY_DN9003_c0_g1 | Light response |
| GBF | GBF1_ARATH | G-box-binding factor 1 | TRINITY_DN2605_c2_g1 | Light response |
| GLK | GLK1_ARATH | Transcription activator GLK1 | TRINITY_DN969_c0_g1 | Light response |
| HAT | HAT1_ARATH | Homeobox-leucine zipper protein HAT1 | TRINITY_DN4311_c1_g1 | Light response |
| HY | | Transcription factor HY5-like | TRINITY_DN34596_c0_g1 | Light response |
| MYC | MYC2_SOLLC | Transcription factor MYC2 | TRINITY_DN4962_c0_g1 | Light response |

| | | | | |
|---|---|---|---|---|
| NAC | | NAC domain-containing protein | TRINITY_DN3383_c0_g1 | Light response |
| PHOT | | Phototropin-2 | TRINITY_DN282_c3_g1 | Light response |
| PHY | PHYA1_TOBAC | Phytochrome A1 | TRINITY_DN2460_c0_g1 | Light response |
| SIG | SIGA_ARATH | RNA polymerase sigma factor sigA | TRINITY_DN4315_c1_g1 | Light response |
| UVR | UVR8_ARATH | Ultraviolet-B receptor UVR8 | TRINITY_DN1267_c1_g1 | Light response |
| CCA | LHY_ARATH | Protein LHY | TRINITY_DN22210_c0_g1 | Circadian rhythm |
| CDF | CDF2_ARATH | Cyclic dof factor 2 | TRINITY_DN3894_c1_g1 | Circadian rhythm |
| CK | CSK21_ARATH | Casein kinase II subunit alpha-1 | TRINITY_DN1064_c0_g2 | Circadian rhythm |
| COL | COL4_ARATH | Zinc finger protein CONSTANS-LIKE 4 | TRINITY_DN3547_c1_g1 | Circadian rhythm |
| COP | COP1_ARATH | E3 ubiquitin-protein ligase COP1 | TRINITY_DN2263_c1_g1 | Circadian rhythm |
| ELF | | Protein EARLY FLOWERING 3 | TRINITY_DN11682_c0_g1 | Circadian rhythm |
| FT | | Glutaminyl-peptide cyclotransferase | TRINITY_DN17372_c0_g1 | Circadian rhythm |
| GI | GIGAN_ARATH | Protein GIGANTEA | TRINITY_DN8078_c0_g1 | Circadian rhythm |
| LHY | LHY_ARATH | Protein LHY | TRINITY_DN7854_c1_g1 | Circadian rhythm |
| LUX | MYBC1_ARATH | Transcription factor MYBC1 | TRINITY_DN1624_c0_g1 | Circadian rhythm |
| MYB | MYB4_ARATH | Transcription repressor MYB4 | TRINITY_DN2554_c0_g1 | Circadian rhythm |
| RVE | RVE7L_ARATH | Protein REVEILLE 7-like | TRINITY_DN10821_c0_g1 | Circadian rhythm |
| SPA | SPA1_ARATH | Protein SUPPRESSOR OF PHYA-105 1 | TRINITY_DN1322_c1_g1 | Circadian rhythm |
| TCP | PIP25_ARATH | Probable aquaporin PIP2-5 | TRINITY_DN9818_c0_g1 | Circadian rhythm |

**Table S4**. **(related with fig S4**)

Annotation of selected genes differentially expressed across experimental variables without a known role in CCM.

| **Uniprot_gene** | **unigenes** | **Description** |
|---|---|---|
| ATHB7 | TRINITY_DN2613_c0_g1 | Homeobox-leucine zipper protein ATHB-7 |
| BH062 | TRINITY_DN360_c1_g1 | Light response |
| BHLH | TRINITY_DN7311_c0_g1 | Light response |
| CIPK5 | TRINITY_DN1514_c1_g1 | CBL-interacting serine/threonine-protein kinase 5 |
| CIPKB | TRINITY_DN21721_c0_g1 | CBL-interacting serine/threonine-protein kinase 11 |
| EF1A | TRINITY_DN8030_c3_g1 | Elongation factor 1-alpha |
| EFMT4 | TRINITY_DN2747_c0_g1 | EEF1A lysine methyltransferase 4 |
| EPS1 | TRINITY_DN2901_c0_g1 | Protein ENHANCED PSEUDOMONAS SUSCEPTIBILITY 1 |
| EXPA2 | TRINITY_DN162_c0_g2 | Expansin-A2 |
| EXTN3 | TRINITY_DN7986_c0_g1 | Extensin-3 |
| HDT1 | TRINITY_DN4285_c1_g2 | Histone deacetylase HDT1 |
| HSP16 | TRINITY_DN68667_c0_g1 | 18.5 kDa class I heat shock protein |
| HYBA1 | TRINITY_DN1856_c0_g1 | Non-reducing end beta-L-arabinofuranosidase |
| KAI2 | TRINITY_DN7983_c1_g1 | Probable esterase KAI2 |
| LSH3 | TRINITY_DN12674_c0_g1 | Protein LIGHT-DEPENDENT SHORT HYPOCOTYLS 3 |
| LSH6 | TRINITY_DN12674_c0_g1 | Protein LIGHT-DEPENDENT SHORT HYPOCOTYLS 6 |
| MYB4 | TRINITY_DN2554_c0_g1 | Transcription repressor MYB4 |
| NAP2 | TRINITY_DN7132_c0_g5 | NAC domain-containing protein 2 |
| NCED1 | TRINITY_DN2687_c0_g2 | 9-cis-epoxycarotenoid dioxygenase NCED1, chloroplastic |
| NFD4 | TRINITY_DN13340_c0_g1 | Protein NUCLEAR FUSION DEFECTIVE 4 |
| NIA | TRINITY_DN6723_c1_g1 | Nitrate reductase [NADH] |
| NLTP | TRINITY_DN79002_c1_g1 | Probable non-specific lipid-transfer protein AKCS9 |
| PAR1 | TRINITY_DN1658_c0_g1 | Phenylacetaldehyde reductase |
| PER2 | TRINITY_DN11931_c0_g1 | Cationic peroxidase 2 |
| PGLR | TRINITY_DN2996_c0_g1 | Probable polygalacturonase |
| PUMP4 | TRINITY_DN12670_c1_g1 | Mitochondrial uncoupling protein 4 |
| PURA2 | TRINITY_DN2249_c0_g2 | Adenylosuccinate synthetase 2, chloroplastic |
| RL261 | TRINITY_DN8213_c0_g1 | 60S ribosomal protein L26-1 |
| RL9 | TRINITY_DN56917_c0_g1 | 60S ribosomal protein L9 |
| RS3A | TRINITY_DN11338_c0_g2 | 40S ribosomal protein S3a |
| SCL15 | TRINITY_DN19737_c0_g1 | Scarecrow-like protein 15 |
| VPE1 | TRINITY_DN9527_c0_g1 | Vacuolar-processing enzyme |
| WRKY7 | TRINITY_DN13587_c0_g1 | Probable WRKY transcription factor 7 |
| WTR32 | TRINITY_DN5341_c0_g2 | WAT1-related protein At4g08300 |

**Table S5. (related with visium spatial gene expression results)**

Sequencing and reads mapping statistics across Visium mRNA libraries.

| Sample ID | n_spots | m_ Genes | n_reads | m_reads _per_spot | n_reads_ map_confidently | Fraction Reads in Spots Under Tissue | Total Genes Detected | Median UMI Counts per Spot | Fraction of Spots Under Tissue |
|---|---|---|---|---|---|---|---|---|---|
| S1_A1_PO1_7h_WW | 1426 | 53 | 150099218 | 105258.92 | 0.04 | 0.92 | 138 | 368.5 | 0.29 |
| S2_B1_PO1_23h_WW | 1972 | 57 | 199760709 | 101298.53 | 0.03 | 0.98 | 139 | 352.5 | 0.40 |
| S3_D1_PO1_23h_D | 1751 | 26 | 121129683 | 69177.43 | 0.01 | 0.80 | 136 | 44 | 0.35 |
| S4_A1_PO1_7h_WW | 1925 | 40 | 192674632 | 100090.72 | 0.03 | 0.90 | 137 | 167 | 0.39 |
| S5_B1_PO1_23h_WW | 3757 | 34 | 322216248 | 85764.24 | 0.02 | 0.96 | 139 | 95 | 0.75 |
| S6_D1_PO1_23h_D | 1575 | 25 | 117371115 | 74521.34 | 0.01 | 0.75 | 135 | 41 | 0.32 |

**Table S6**. **(related with figures 6-7)**

Sensitivity analysis of pFBA results of major reactions for Rubisco carboxylation to oxygenation ratio ($V_c/V_o$) for $C_4$ in bundle sheath.

| $V_c/V_o$ | 10 | 20 | 40 | 80 | 200 |
|---|---|---|---|---|---|
| Phloem output | 3.14 | 3.32 | 3.43 | 3.48 | 3.51 |
| PEP carboxylation_M_day | 70.00 | 70.00 | 70.00 | 70.00 | 70.00 |
| PEP carboxylation_BS_day | 0 | 0 | 0 | 0 | 0 |
| PEP carboxylation_M_night | 47.23 | 54.48 | 58.23 | 60.15 | 61.31 |
| PEP carboxylation_BS_night | 8.68 | 8.64 | 8.66 | 8.67 | 8.67 |
| Rubsico carboxylation_M_day | 0 | 0 | 0 | 0 | 0 |
| Rubsico carboxylation_BS_day | 133.51 | 137.53 | 136.63 | 140.71 | 141.36 |
| Rubsico carboxylation_M_night | 0 | 0 | 0 | 0 | 0 |
| Rubsico carboxylation_BS_night | 0 | 0 | 0 | 0 | 0 |
| CO2_M_day | 70.00 | 70.00 | 70.00 | 70.00 | 70.00 |
| CO2_BS_day | 0 | 0 | 0 | 0 | 0 |
| CO2_M_night | 40.79 | 47.44 | 50.91 | 52.69 | 53.77 |
| CO2_BS_night | 0 | 0 | 0 | 0 | 0 |

**Table S7**. **(related with figures 6-7)**

pFBA results for major reactions of additional modelling scenarios in mesophyll (M) or bundle sheath (BS) at daytime (day) and night time (night). Major reactions are Phloem output, PEP carboxylation (PEP), Rubisco carboxylation (Rubisco) and $CO_2$ absorption ($CO_2$). All the scenarios are modelling under the drought condition. Scenario 1): blocking malate transfer between mesophyll and bundle sheath (bMalT). Scenario 2):blocking malate storage in mesophyll (bMS), bundle sheath (bBSS), or both (bMBSS). Scenario 3): CAM with $C_3$ or $C_4$anatomy. C3+CAM: both $C_3$and CAM activity (both daytime and night time $CO_2$ uptake) allowedwith a $C_3$ anatomy ($CO_2$directly diffuses into mesophyll, bundle sheath is considered an inner mesophyll with a longer distance to stomata); CAM_C3: only CAM (nighttime $CO_2$ uptake) occurswith $C_3$anatomy ; CAM_C4: CAM process (nighttime $CO_2$ uptake) with $C_4$anatomy ($CO_2$ can not directly diffuse into bundle sheath); C4+CAM_C4: C4 and CAM (both daytime and night time $CO_2$ uptake) can occurwith $C_4$anatomy ($CO_2$ can not directly diffuse into bundle sheath), which can be used as the reference for all the above scenarios.

| SCENARIO | bMalT | bMS | bBSS | bMBSS | C3+CAM_ C3 | CAM_C3 | CAM_C4 | C4+CAM_ C4 |
|---|---|---|---|---|---|---|---|---|
| **Phloem output** | 3.14 | 3.14 | 3.14 | 2.69 | 2.56 | 2.48 | 2.79 | 3.14 |
| **PEP_M_day** | 70.00 | 70.00 | 70.00 | 70.00 | 0 | 0 | 0 | 70.00 |
| **PEP_BS_day** | 0 | 0 | 0 | 0 | 0 | 0 | 0 | 0 |
| **PEP_M_night** | 47.49 | 49.53 | 50.72 | 24.74 | 29.91 | 77.05 | 108.65 | 47.23 |
| **PEP_BS_night** | 8.42 | 6.38 | 5.19 | 6.50 | 4.28 | 29.99 | 9.76 | 8.68 |
| **Rubisco_M_day** | 0 | 0 | 0 | 0 | 88.94 | 85.33 | 0 | 0 |
| **Rubisco_BS_day** | 133.51 | 133.51 | 133.51 | 164.45 | 36.40 | 33.49 | 125.49 | 133.51 |
| **Rubisco_M_night** | 0 | 0 | 0 | 0 | 0 | 0 | 0 | 0 |
| **Rubisco _BS_night** | 0 | 0 | 0 | 23.32 | 0 | 0 | 0 | 0 |
| **CO2_M_day** | 70.00 | 70.00 | 70.00 | 70 | 40.00 | 0 | 0 | 70.00 |
| **CO2_M_night** | 40.79 | 40.79 | 40.79 | 24.74 | 30.00 | 0 | 0 | 0 |

### Online Supplementary Materials

**Captions for Table S8 (related to 'LMD-RNA Sequencing and read alignments' results).**

Estimated counts of reads mapped to each unigene per LMD library.

**Captions for Table S9 (related to 'Global transcriptional differences across experimental variables' results).**

Transcriptome-wide gene annotation and differential expression statistics.

**Captions for Table S10 (related to 'Transcriptional changes in CCM-related genes across cell types' results)**

CCM-related (or hypothetically related) genes. Functional and phylogenetic annotation into gene lineage and/or gene families. Differential expression statistics.

**Captions for Table S11 (related to 'Transcriptional changes in CCM-related genes across cell types' results, and to figs. 3-4-5-S2-S3).**

Representative contigs (highest expressed) by gene lineage and/or gene family. Differential expression statistics.

**Captions for Table S12 (related to 'Flux balance model' results and to Figs. 6-7-S6-S7)**

FBA results summary.

**Captions for Data S1 to S6 (related to 'Visium gene expression' results)**
Output files quantification and analysis of transcripts abundance across Visium samples. Files can be visualized using the software Loupe Browser (10X Genomics; https://support.10xgenomics.com/single-cell-gene-expression/software/visualization/latest/what-is-loupecell-browser). The list of gene features to display in Loupe are found in table S1
